# Impact of rare and common genetic variants on diabetes diagnosis by hemoglobin A1c in multi-ancestry cohorts: The Trans-Omics for Precision Medicine Program

**DOI:** 10.1101/643932

**Authors:** Chloé Sarnowski, Aaron Leong, Laura M Raffield, Peitao Wu, Paul S de Vries, Daniel DiCorpo, Xiuqing Guo, Huichun Xu, Yongmei Liu, Xiuwen Zheng, Yao Hu, Jennifer A Brody, Mark O Goodarzi, Bertha A Hidalgo, Heather M Highland, Deepti Jain, Ching-Ti Liu, Rakhi P Naik, James A Perry, Bianca C Porneala, Elizabeth Selvin, Jennifer Wessel, Bruce M Psaty, Joanne E Curran, Juan M Peralta, John Blangero, Charles Kooperberg, Rasika Mathias, Andrew D Johnson, Alexander P Reiner, Braxton D Mitchell, L Adrienne Cupples, Ramachandran S Vasan, Adolfo Correa, Alanna C Morrison, Eric Boerwinkle, Jerome I Rotter, Stephen S Rich, Alisa K Manning, Josée Dupuis, James B Meigs, on behalf of the Trans-Omics for Precision Medicine (TOPMed) Diabetes and TOPMed Hematology and Hemostasis working groups and the NHLBI TOPMed Consortium

## Abstract

Hemoglobin A1c (HbA1c) is widely used to diagnose diabetes and assess glycemic control in patients with diabetes. However, nonglycemic determinants, including genetic variation, may influence how accurately HbA1c reflects underlying glycemia. Analyzing the NHLBI Trans-Omics for Precision Medicine (TOPMed) sequence data in 10,338 individuals from five studies and four ancestries (6,158 Europeans, 3,123 African-Americans, 650 Hispanics and 407 East Asians), we confirmed five regions associated with HbA1c (*GCK* in Europeans and African-Americans, *HK1* in Europeans and Hispanics, *FN3K/FN3KRP* in Europeans and *G6PD* in African-Americans and Hispanics) and discovered a new African-ancestry specific low-frequency variant (rs1039215 in *HBG2/HBE1*, minor allele frequency (MAF)=0.03). The most associated *G6PD* variant (p.Val98Met, rs1050828-T, MAF=12% in African-Americans, MAF=2% in Hispanics) lowered HbA1c (−0.88% in hemizygous males, −0.34% in heterozygous females) and explained 23% of HbA1c variance in African-Americans and 4% in Hispanics. Additionally, we identified a rare distinct *G6PD* coding variant (rs76723693 - p.Leu353Pro, MAF=0.5%; −0.98% in hemizygous males, −0.46% in heterozygous females) and detected significant association with HbA1c when aggregating rare missense variants in *G6PD*. We observed similar magnitude and direction of effects for rs1039215 (*HBG2*) and rs76723693 (*G6PD*) in the two largest TOPMed African-American cohorts and replicated the rs76723693 association in the UK Biobank African-ancestry participants. These variants in *G6PD* and *HBG2* were monomorphic in the European and Asian samples. African or Hispanic ancestry individuals carrying *G6PD* variants may be underdiagnosed for diabetes when screened with HbA1c. Thus, assessment of these variants should be considered for incorporation into precision medicine approaches for diabetes diagnosis.

## Introduction

Hemoglobin A1c (HbA1c) is a convenient indirect measure of long-term exposure to blood glucose concentrations. HbA1c estimates the proportion of glycated hemoglobin in the blood, an irreversible chemical modification of the hemoglobin molecule by blood glucose.^1^ As HbA1c reflects average ambient glycemia over the previous 2-3 months, the life of an erythrocyte, it is a commonly used as a test to diagnose diabetes (MIM: 125853 and 222100) and estimate glycemic control in patients with diabetes. However, non-glycemic variation in HbA1c due to differences in erythrocyte turnover can influence how accurately HbA1c reflects underlying glycemia.^2^

We previously conducted a trans-ethnic genome-wide association study (GWAS) meta-analysis of HbA1c in 159,940 individuals from four ancestries (European, African, East Asian, and South Asian). We identified 60 common (minor allele frequency, MAF, greater than 5%) genetic variants associated with HbA1c, of which 19 were classified as ‘glycemic’ and 22 as ‘erythrocytic’ based on the probable biological mechanism through which they appeared to influence HbA1c levels.^3^ Genetic variants affecting HbA1c via erythrocyte biological pathways may lead to diagnostic misclassification of ambient glycemia and thus diabetes status. HbA1c GWAS have so far focused on genetic variants imputed to HapMap,^4^ therefore genetic discovery efforts have been focused on common variants. Low-frequency (0.5% < MAF < 5%) and rare (MAF < 0.5%) genetic variants and their associated impact on the diagnostic accuracy of HbA1c have not been systematically examined but are suspected to occur.^5, 6^ Further, previous GWAS have shown that the combined effect of HbA1c-related common variants causes differences in HbA1c that were three times greater in individuals of African ancestry compared with those of European ancestry [0.8 (%-units) vs. 0.25 (%-units)].^3^ This relatively large difference in HbA1c was mainly driven by a single African-ancestry specific missense variant (p.Val98Met, rs1050828-T) in the Glucose-6-phosphatase Dehydrogenase gene (*G6PD* [MIM:305900]), which causes G6PD deficiency (MIM: 300908).^7, 8^ As the genetic architecture of HbA1c appears to differ by ancestry, as does type 2 diabetes risk, it is imperative to understand the genetic basis of HbA1c in different ancestral groups to ensure that large-effect ancestry-specific variants, similar to the *G6PD* variant, are uncovered.^9^

Here, we sought to confirm common and identify low frequency and rare genetic variants associated with HbA1c through association analyses in diabetes-free individuals from four ancestries using whole genome sequencing (WGS) data from the NIH National Heart, Lung, and Blood Institute (NHLBI) Trans-Omics for Precision Medicine (TOPMed) program. We hypothesized that variants in the low frequency spectrum with relatively large effects will be detected even with modest sample sizes. To uncover additional distinct erythrocytic variants associated with HbA1c, we performed fine-mapping using sequential conditional association analyses of erythrocytic loci that reached genome-wide significance in this study as well as gene-based tests.

## Methods

### Populations and participants

We included in our analyses 10,338 TOPMed participants without diabetes from five cohorts: the Old Order Amish study (N=151), the Atherosclerosis Risk in Communities Study (ARIC, N=2,415), the Framingham Heart Study (FHS, N=2,236), the Jackson Heart Study (JHS, N=2,356) and the Multi-Ethnic Study of Atherosclerosis (MESA, N=3,180) representing four ancestry groups: Europeans (EA, N=6,158), African-Americans (AA, N=3,123), Hispanics/Latinos (HA, N=650) and East Asians (AS, N=407) (**Supplementary Table 1**). Descriptions of each cohort are available in the **Supplementary Text**. Diabetes was defined as fasting glucose (FG) ≥ 7 mmol/L after ≥ 8 hours, HbA1c ≥ 6.5%-units, 2-hour glucose by an oral glucose tolerance test ≥ 11.1 mmol/L, non-fasting glucose ≥ 11.1 mmol/L, physician diagnosed diabetes, self-reported diabetes, or use of an antidiabetic medication. Measures of FG and 2-hour glucose made in whole blood were corrected to plasma levels using the correction factor of 1.13. Individual studies applied further sample exclusions where applicable, including pregnancy and type 1 diabetes status.

### Measurement of HbA1c and erythrocytic traits

The National Glycohemoglobin Standardization Program (NGSP) certified assays^10^ used to measure HbA1c in each cohort are indicated in **Supplementary Table 2**. HbA1c was expressed in NGSP %-units. Measurements for red blood cell (RBC) count (×1012/L), hemoglobin (HB; g/dL), hematocrit (HCT; %), mean corpuscular volume (MCV; fL), mean corpuscular hemoglobin (MCH; pg), mean corpuscular hemoglobin concentration (MCHC; g/dL), and red blood cell distribution width (RDW; %) were obtained from complete blood count panels performed using standard assays.

### Whole genome sequencing

The NHLBI TOPMed program provided WGS, performed at an average depth of 38× by several sequencing centers (New York Genome Center; Broad Institute of MIT and Harvard; University of Washington Northwest Genomics Center; Illumina Genomic Services; Macrogen Corp.; and Baylor Human Genome Sequencing Center), using DNA from blood. Details regarding the laboratory methods, data processing and quality control are described on the TOPMed website (see URL in the web resources section) and in documents included in each TOPMed accession released on the database of Genotypes and Phenotypes (dbGaP). Processing of whole genome sequences was harmonized across genomic centers using a standard pipeline.^11^

We used TOPMed ‘freeze 5b’ that comprised 54,508 samples. Variant discovery and genotype calling were performed jointly across TOPMed parent studies for all samples using the GotCloud pipeline. A support vector machine quality filter was trained using known variants (positive training set) and Mendelian-inconsistent variants (negative training set). The TOPMed data coordinating center performed additional quality control checks for sample identity issues including pedigree errors, sex discrepancies, and genotyping concordance. After site level filtering, TOPMed freeze 5b consisted of ~438 million single nucleotide variants (SNVs) and ~33 million short insertion-deletion variants. Read mapping was done using the 1000 Genomes Project reference sequence versions for human genome build GRCh38.

The study was approved by the appropriate institutional review boards (IRB) and informed consent was obtained from all participants.

### Statistical analyses

In each ancestry, we performed pooled WGS association analysis of HbA1c using Genesis on the Analysis Commons (see URL in the web resources section).^12^ We used linear mixed effect models to test the association of HbA1c with the SNVs individually while adjusting for sex, age at HbA1c measurement and study, allowing for heterogeneous variance across study groups. We accounted for relatedness using an empirical kinship matrix (Genetic Relationship Matrix, GRM). We excluded from our single-SNV analyses variants with a minor allele count (MAC) less than 20 across the combined samples. We used a significance threshold of P < 2×10^-8^ to report an association as genome-wide significant for common, low frequency and rare genetic variants, which was slightly more stringent than the widely adopted P-value threshold of 5×10^-8^ in GWAS, based on estimations for genome-wide significance for WGS studies in UK10K.^13, 14^ For the X chromosome, genotypes were coded as 0 and 2 for males and 0, 1 and 2 for females, and sex-stratified analyses (analyses conducted separately in males and females) were also performed. We also performed additional analyses in AA, adjusted on the sickle cell trait (SCT) variant rs334, as well as haplotype analyses with this variant, using the R package haplo.stats.

Because loci that influence HbA1c through erythrocytic mechanisms are expected to cause nonglycemic variation in HbA1c, we sought to classify HbA1c-associated loci as ‘glycemic’ or ‘erythrocytic’ using mediation analyses on FG, HB, MCV, MCH or MCHC. If the HbA1c-variant association effect size changed by more than 25% upon adding FG to the regression model, the variant was classified as ‘glycemic’. If a variant was not ‘glycemic’, and if the effect size changed by more than 25% upon adding HB, MCV, MCH, or MCHC to the regression model, the variant was classified as ‘erythrocytic’. If association effect sizes were unchanged by mediation adjustment, then the variant was considered ‘unclassified’.

If HbA1c-associated variants remained unclassified by the mediation analyses, we used association analysis results with FG to classify them as ‘glycemic’ (*P* < 0.005), and association analysis results with erythrocytic traits (HB, MCV, MCH, MCHC, RBC, HCT or RDW) to classify them as ‘erythrocytic’ (*P* < 0.005). Association results for FG were obtained from a pooled association analysis performed using a linear mixed effect model in Genesis, adjusted on age, age squared, sex, BMI and self-report ancestry in TOPMed freeze 5b (N=26,883). Association results for erythrocytic traits were obtained from several sources. We first used single variant association analyses performed using a score test in Genesis and adjusted for age, sex, and study, with a GRM, while allowing for heterogeneous variance across study groups, in TOPMed freeze 5b (N=25,080). The number of individuals included in each analysis, per study, is available in **Supplementary Table 3**. Analyses were performed at the University of Washington using the TOPMed pipeline. We then used association results from published GWAS in European (UK Biobank and INTERVAL studies; N=173,480),^15^ Hispanic (Hispanic Community Health Study and Study of Latinos (HCHS/SOL); N=12,502),^16^ and African-American participants (Continental Origins and Genetic Epidemiology [COGENT] Network; N~16,500,^17^ and the Candidate gene Association Resource (CARe) Project N=7,112),^18^ and exome genotyping in 130,273 multi-ethnic individuals.^19^

For loci classified as ‘erythrocytic’, we performed sequential conditional analyses to determine the number of distinct signals in each region. The regions were defined based on linkage disequilibrium (LD) plots. We used a Bonferroni correction for the number of SNVs in the region with MAC ≥ 20 to define a signal as distinct. We also performed gene-based and burden tests in Genesis using different selection of rare genetic variants (MAF ≤ 1%) based on functional annotations (missense, high confidence loss of function or synonymous variants).

Results from each analysis were combined across ancestries or across sex by meta-analysis using METAL.^20^ The heterogeneity test between males and females was calculated using the following formula: 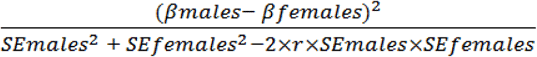. This assessed the difference in effect sizes between males (βmales) and females (βfemales) while accounting for correlation (r) between male and female statistics due to relatedness and was calculated outside of METAL. LD calculations in the TOPMed data and regional plots were done, within ancestry group, using the Omics Analysis, Search and Information System (OASIS, see URL in the web resources section). Functional annotations were performed using the WGS Annotator (WGSA).^21^

We used the UK Biobank (UKBB, (see URL in the web resources section), a prospective cohort study with deep genetic and phenotypic data collected on approximately 500,000 individuals from across the United Kingdom, aged between 40 and 69 at recruitment, as an independent sample for external replication of our findings. The centralized analysis of the genetic data, including genotype quality, population structure and relatedness of the genetic data, and efficient phasing and genotype imputation has been described extensively elsewhere.^22^ Two similar arrays were used for genotyping (Applied Biosystems UK Biobank Lung Exome Variant Evaluation and UK Biobank Axiom Arrays) and pre-phasing was performed using markers present on both arrays. Phasing on the autosomes was carried out using SHAPEIT3 and 1000 Genomes phase 3 panel to help with the phasing of non-European ancestry samples. Imputations were carried out using the IMPUTE4 program with the Haplotype Reference Consortium (HRC) reference panel or with a merged UK10K and 1000 Genomes phase 3 reference panel. For chromosome X, haplotype estimation and genotype imputation were carried out separately on the pseudo-autosomal and non-pseudo autosomal regions.

We identified UKBB African-ancestry participants using the following six self-reported ethnicities: “Caribbean”, “African”, “Black or Black British”, “Any other Black background”, “White and Black African” and “White and Black Caribbean”. We excluded participants with diabetes defined by the use of antidiabetic medication, self-reported physician diagnosis, FG ≥ 7 mmol/L or non-fasting glucose ≥ 11.1 mmol/L.

We generated principal components using smartPCA (see URL in the web resources section)^23, 24^ based on 72,300 common genetic variants in low LD selected with PLINK.^25^ For each PC, ethnic outliers lying more than 6SD away from the mean were excluded. The same genetic variants were used to calculate an empirical kinship matrix using EPACTS (see URL for EPACTS documentation in the web resources section). We used a linear mixed effect model in R to evaluate the association of each variant with HbA1c, adjusting for age, sex, 10 PCs and using the empirical kinship matrix.

## Results

### HbA1c-associated regions using WGS

We included in our analyses 10,338 TOPMed participants representing four ancestry groups: Europeans (EA, N=6,158), African-Americans (AA, N=3,123), Hispanics/Latinos (HA, N=650) and East Asians (AS, N=407; **Supplementary Table 1**). TOPMed studies were composed of middle to older-aged participants of EA, AA, HA or AS ancestry with comparable mean HbA1c, mean fasting glucose (FG) and mean hemoglobin (HB, **Supplementary Table 4**). A total of 13,079,661 variants (EA), 21,443,543 variants (AA), 9,567,498 variants (HA), and 6,567,324 variants (AS) passed filters and were included in the analyses. QQ-plots and Manhattan plots of WGS associations with HbA1c from ancestry-specific analyses and the meta-analysis are provided in Figure 1 **and Supplementary Figures 1-2**. Using a significance threshold of P < 2×10^-8^ to report an association as genome-wide significant, we detected five regions associated with HbA1c, including one novel locus (low-frequency AA-specific variant, rs1039215 in *HBG2* [MIM:142250] */HBE1* [MIM:142100], MAF=0.03), and a sixth locus reaching suggestive evidence (rare AA-specific variant, rs551601853, near *XPNPEP1* [MIM:602443], MAF=0.003, *P* < 5 × 10^-7^). Regional plots for the *HBG2/HBE1* and the *XPNPEP1* loci are provided in **Supplementary Figures 3 and 4**. The four other single nucleotide variants (SNVs) were located in regions previously identified as associated with HbA1c in trans-ethnic meta-analyses^3^: rs2971670 in *GCK* on chromosome 7 (P_meta_=1.7 10^-9^, mainly associated in EA and AA, r^2^=0.59, D’=1 with rs4607517, the index SNV in published GWAS), rs17476364 in *HK1* [MIM:142600] on chromosome 10 (P_meta_=3.1 × 10^-21^, mainly associated in EA and HA, r^2^=0.10, D’=1 with rs10823343, r^2^=0.18, D’=0.50 with rs4745982, index SNVs in published GWAS), rs113373052 in *FN3K* [MIM:608425] */FN3KRP* [MIM:611683] on chromosome 17 (P_meta_=4.5 × 10^-10^, associated in EA, r^2^=0.92, D’=0.99, with rs1046896, index SNV in published GWAS) and rs1050828 in *G6PD* on chromosome X (P_meta_=5.1 × 10^-210^, associated in AA and HA, monomorphic in the other ancestries)^3^. The top SNV in each region detected at the genome-wide threshold (P < 2 × 10^-8^) is indicated in Table 1 and at the sub-genome-wide threshold (P < 5 × 10^-7^) in **Supplementary Table 5**.

**Figure 1:**
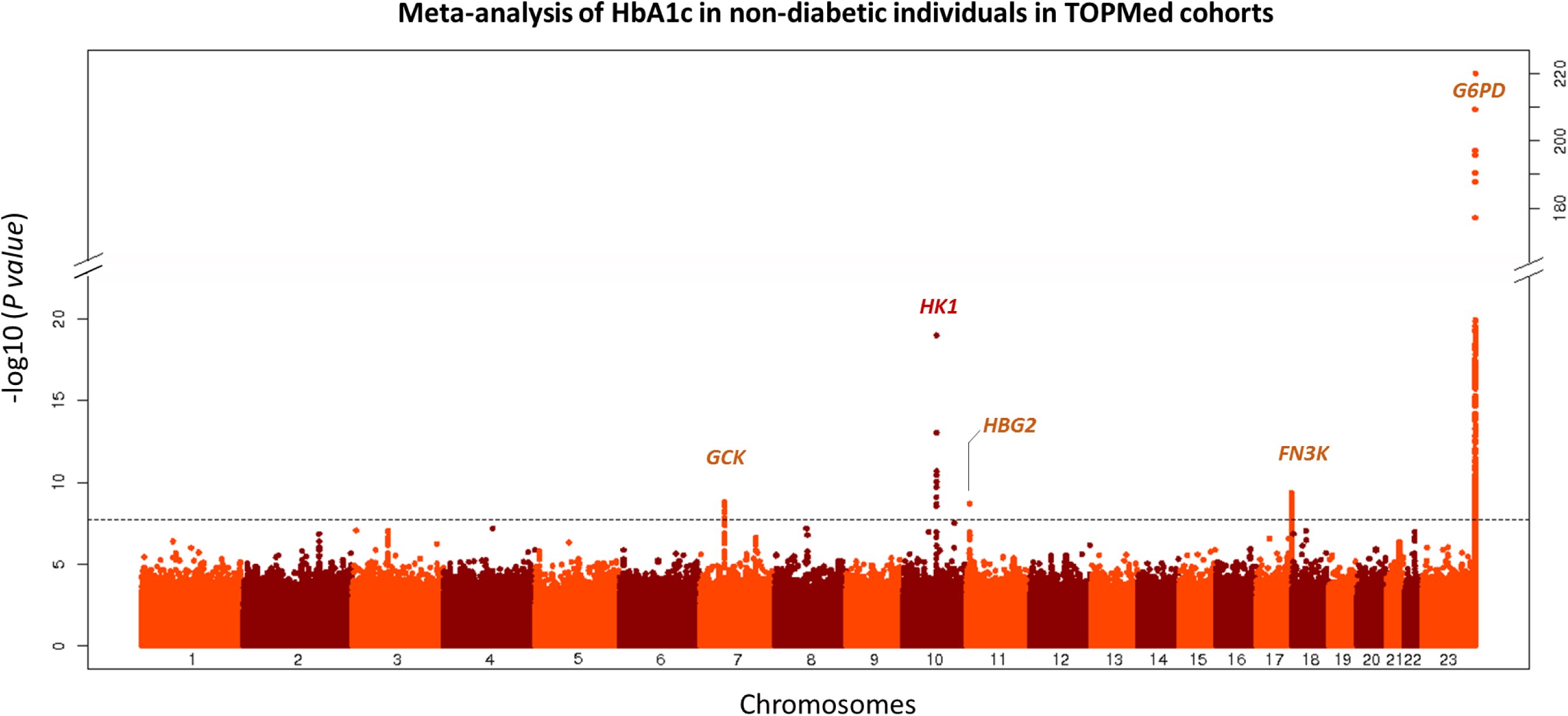
Manhattan-plots of the meta-analysis of HbA1c in non-diabetic individuals in TOPMed cohorts. The –log10(P-value) for each single nucleotide variant on the y-axis is plotted against the build 38 genomic position on the x-axis (chromosomal coordinate). The dashed horizontal line indicates the genome-wide significance threshold of *P* = 2×10^-8^. The y-axis was truncated for ease of interpretation.

**Table 1:** Most associated single nucleotide variants in the regions detected at the genome-wide level (P<2×10^-8^) by ancestry and in the meta-analysis of HbA1c in non-diabetic individuals in TOPMed cohorts

|  |  |  |  | Meta-analysis |  |  | Europeans (N=6,158) |  |  |  | African-Americans (N=3,123) |  |  |  | Hispanics (N=650) |  |  |  | Asians (N=407) |  |  |  |
| --- | --- | --- | --- | --- | --- | --- | --- | --- | --- | --- | --- | --- | --- | --- | --- | --- | --- | --- | --- | --- | --- | --- |
| MarkerID<sup>b</sup> | rsID | Closest Gene | Ref/Alt Allele | Beta | SE | *P value* | MAF<sup>a</sup> | Beta | SE | *P value* | MAF<sup>a</sup> | Beta | SE | *P value* | MAF<sup>a</sup> | Beta | SE | *P value* | MAF<sup>a</sup> | Beta | SE | *P value* |
| 7:44186502 | rs2971670 | *GCK* | C/T | 0.04 | 0.01 | 1.7E-09 | 0.18 | 0.03 | 0.01 | 1.6E-05 | 0.18 | 0.06 | 0.01 | 1.6E-04 | 0.21 | 0.05 | 0.03 | 0.04 | 0.19 | 0.04 | 0.03 | 0.20 |
| 10:69334748 | rs17476364 | *HK1* | T/C | -0.09 | 0.01 | 3.1E-21 | 0.10 | -0.08 | 0.01 | 3.2E-18 | 0.02 | -0.09 | 0.04 | 0.04 | 0.06 | -0.15 | 0.04 | 5.4E-04 |  |  |  |  |
| 11:5464062 | rs1039215 | *HBG2/*<br>*HBE1* | A/G | -0.21 | 0.04 | 2.0E-09 |  |  |  |  | 0.03 | -0.21 | 0.04 | 2.0E-09 |  |  |  |  |  |  |  |  |
| 17:82739582 | rs113373052 | *FN3K* | C/T | 0.03 | 0.01 | 4.5E-10 | 0.32 | 0.03 | 0.01 | 4.0E-08 | 0.29 | 0.02 | 0.01 | 0.06 | 0.39 | 0.03 | 0.02 | 0.15 | 0.49 | 0.05 | 0.02 | 0.04 |
| X:154536002 | rs1050828 | *G6PD* | C/T | -0.41 | 0.01 | 5.1E-210 |  |  |  |  | 0.12 | -0.41 | 0.01 | 8.4E-205 | 0.02 | -0.37 | 0.07 | 8.3E-07 |  |  |  |  |

<sup>a</sup>MAF: minor allele frequency
<sup>b</sup>MarkerID is defined as Chromosome:Position with positions on Build 38.

### Classification of HbA1c-associated loci by their biological pathways

Because loci that influence HbA1c through erythrocytic pathways are expected to cause nonglycemic variation in HbA1c, we sought to classify HbA1c-associated loci as ‘glycemic’ or ‘erythrocytic’ using mediation analyses and association analyses with glycemic and erythrocytic traits. Mediation analyses and look-up in WGS analysis of FG in TOPMed classified the *GCK* [MIM:138079] variants as glycemic (**Supplementary Tables 6, 7 & 8**). Association analyses with erythrocytic traits in published GWAS showed that rs17476364-C allele, in the *HK1* gene, was positively associated with HB, mean corpuscular volume (MCV), mean corpuscular hemoglobin (MCH), mean corpuscular hemoglobin concentration (MCHC), red blood cell count (RBC), hematocrit (HCT) and red blood cell distribution width (RDW) in EA^15^ and that rs1050828-T allele, in the *G6PD* gene, was positively associated with MCH in AA and MCV in AA and HA and negatively associated with HCT and HB in AA, and RBC and RDW in AA and HA^16–18^ (**Supplementary Tables 6, 8 and 9**). Results of the mediation and association analyses are available in **Supplementary Tables 6, 8 and 9** for the genome-wide variants and **Supplementary Tables 7 and 10** for the sub-genome variants. Among the 26 variants meeting the sub-genome-wide threshold, four were common and 22 were low-frequency or rare. Two of the common HbA1c-associated variants were reported previously:^3^ the glycemic variant at *G6PC2* [MIM:608058] and the erythrocytic variant at *TMPRSS6* [MIM:609862]. Among the genome-wide significant loci, *HK1* (EA) and *G6PD* (AA and HA) were classified as ‘erythrocytic’, as was the low frequency variant rs1039215 in *HGB2/HBE1*, negatively associated with HB, HCT, MCV and MCH. The significant *FN3K/FN3KRP* and *XPNPEP1* variants were unclassified.

### Characterization of distinct signals at erythrocytic HbA1c-associated loci

For loci classified as ‘erythrocytic’ (*HK1*, *HBG2/HBE1* and *G6PD*), we performed sequential conditional analyses to detect distinct signals in addition to the top SNV. The regions were defined as +/- 60kb region around *HK1*, +/- 250kb region around *HBG2/HBE1* and +/- 500kb region around *G6PD*, based on Linkage Disequilibrium (LD) plots (**Supplementary Figures 3, 5 and 6**). We did not detect secondary associations in the *HK1* and *HBG2/HBE1* regions at a threshold of 4.3 × 10^-5^ and 8.5 × 10^-6^ respectively. Interestingly, in the *G6PD* region, the top SNV (rs1050828, p.Val98Met, T-allele frequency 12% in AA, 2% in HA, 0% in EA and AS) was associated with lower HbA1c in both AA and HA. This SNV accounted for 23% of HbA1c variance in AA and 4% in HA. The LD in the *G6PD* region is complex, with a strong haplotype effect in AA and HA (Figure 2). By performing conditional analyses on the top SNV (rs1050828) we were able to detect an additional rare signal (G-allele frequency 0.5%) in AA (rs76723693, p.Leu353Pro, B_cond_=-0.50, P_cond_=2.8 10^-15^) that was distinct from rs1050828 (r2=0.0006, D’=1, Figure 3). The threshold to detect an additional signal was fixed to 1.7×10^-5^ in AA and 4.5×10^-5^ in HA. The three other SNVs located in the same region that were sub-genome-wide significant in the main analysis (rs143745197, rs184539426 and rs189305788) were not significant after adjusting for both rs1050828 and rs76723693. We detected significant or suggestive associations using gene-based (P=9.7 × 10^-11^) and burden tests (P=4.7 × 10^-5^) when aggregating 15 missense rare variants with MAF ≤ 1% in *G6PD* (**Supplementary Table 11**). As association from SKAT was more significant than the association of rs76723693 from single-SNV association analysis, it suggested that several rare missense variants in *G6PD* were associated with HbA1c. We performed additional association and conditional analyses in *G6PD* for missense variants with 10 < minor allele count < 20. In addition to rs76723693, one rare missense variant (rs5030872, MAF=0.002) was suggestively associated with HbA1c (B=-0.64, P=3.4 × 10^-6^) and became significantly associated with HbA1c when adjusting for both rs1050828 and rs76723693 (B_cond_=-0.71, P_cond_=1.3× 10^-9^).

**Figure 2:**
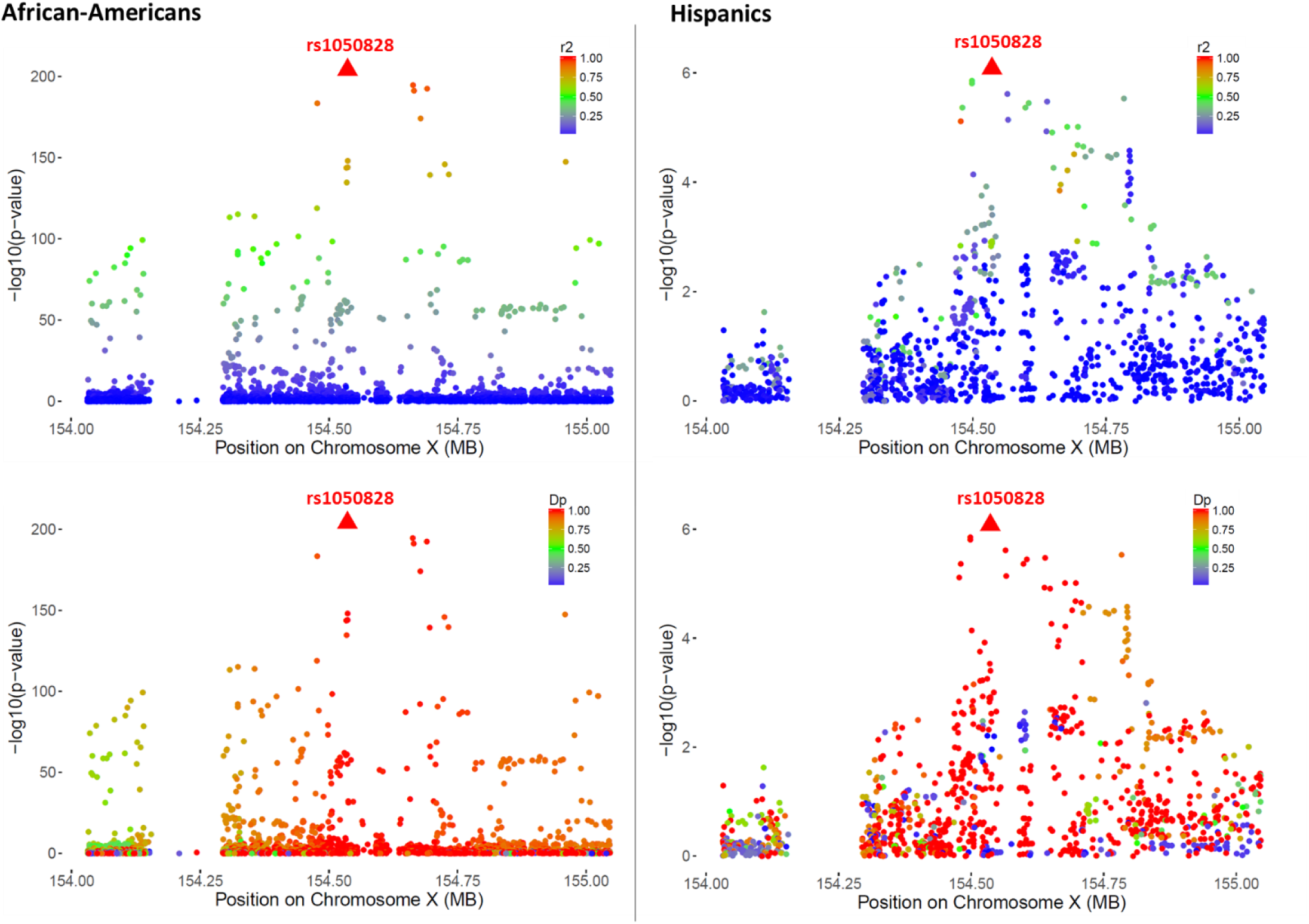
Regional HbA1c association plots in the *G6PD* region in non-diabetic African-Americans and Hispanics in TOPMed cohorts. Association plots are displayed separately in African-Americans (left-side) and in Hispanics (right-side). Single nucleotide variants are plotted with their P-values (-log10 values, y-axis) as a function of build 38 genomic position on chromosome X (x-axis). The local linkage disequilibrium (LD) structure (r2 - top or D’ - bottom) between the top associated single nucleotide variant rs1050828 (red triangle) and correlated proxies is indicated in color according to a blue to red scale from r2/D’=0 to 1 and was calculated separately in non-diabetic African-Americans and Hispanics TOPMed cohorts.

**Figure 3:**
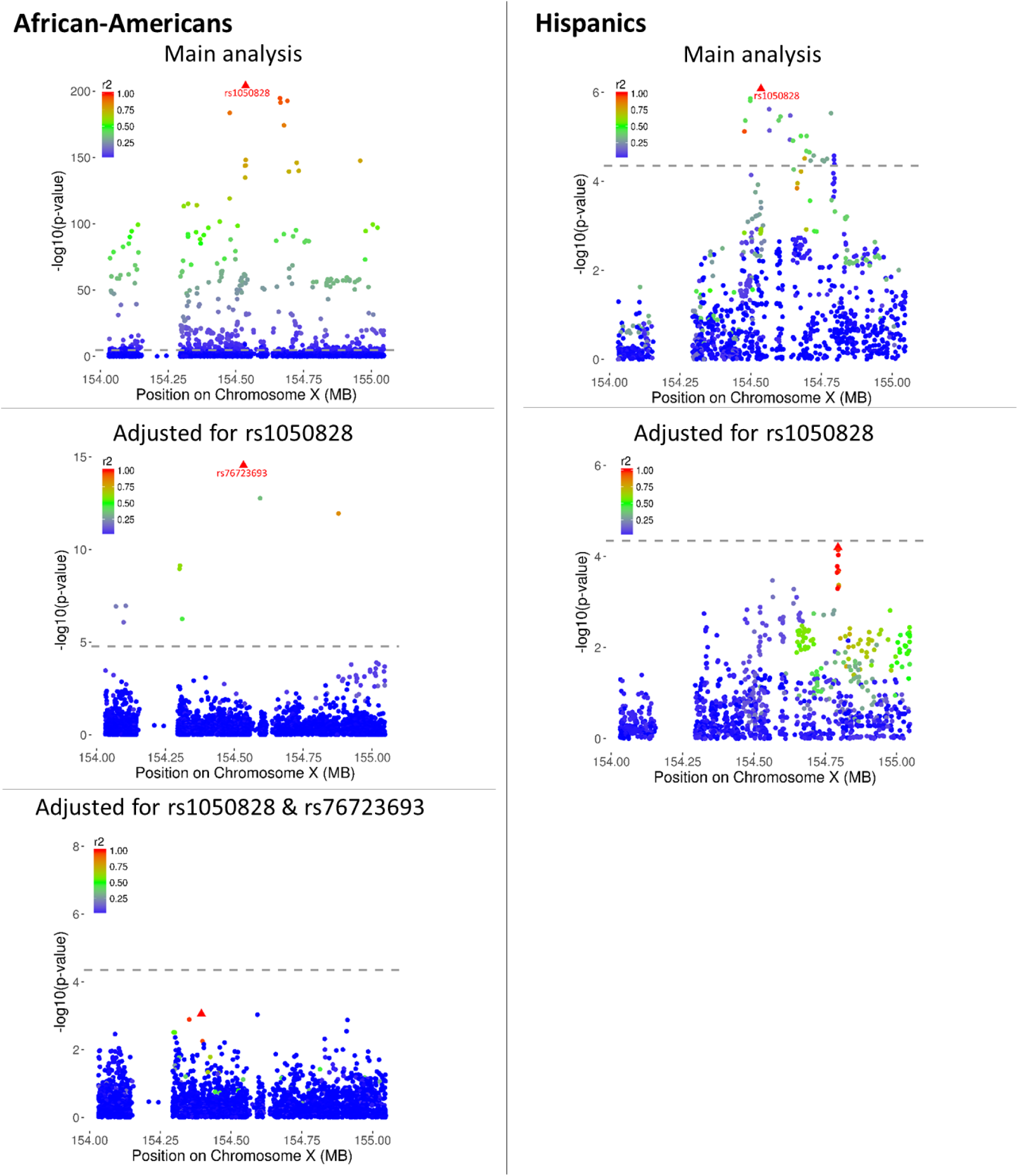
Sequential conditional analyses in the *G6PD* region in non-diabetic African-Americans and Hispanics in TOPMed cohorts. Association plots are displayed separately in African-Americans (left-side) and in Hispanics (right-side). Single nucleotide variants are plotted with their P-values (-log10 values, y-axis) as a function of build 38 genomic position on chromosome X (x-axis). The local linkage disequilibrium (LD) structure (r2 - top or D’ - bottom) between the top associated single nucleotide variant (red triangle) and correlated proxies is indicated in color according to a blue to red scale from r2/D’=0 to 1 and was calculated separately in non-diabetic African-Americans and Hispanics TOPMed cohorts. Significance thresholds to declare an association signal as distinct are indicated by a red line (*P*=1.7 10^-5^ in African-Americans and *P*=4.5×10^-5^ in Hispanics, based on the number of single nucleotide variants in the region with a minor allele count greater than 20).

The *HBG2/HBE1* new variant rs1039215 (P=2.0 × 10^-9^) is located at 327 kb from the SCT variant rs334 (P=2.9 × 10^-8^). A recent paper reported rs1039215 to be in LD (r^2^>0.2) with the SCT variant (rs334, r^2^=0.24, D’=-0.83) and correlated with *HBG2* gene expression in whole blood.^26^ To determine if rs1039215 was distinct from rs334, we calculated LD and performed conditional analyses. In TOPMed, rs1039215 was in modest LD with rs334 (r^2^=0.30, D’=0.88, **Supplementary Figure 3**) and conditioning on rs334 attenuated its association (decrease in p-value from 2.0 × 10^-9^ to 10^-3^, **Supplementary Table 12**), suggesting that the association of rs1039215 with HbA1c may be explained by the SCT rs334 variant. Nevertheless, the haplotype rs334-A / rs1039215-G was more significantly associated with HbA1c than the haplotypes rs334-A / rs1039215-A and rs334-T / rs1039215-G (**Supplementary Table 13**). Thus, the association of the new variant rs1039215 in *HBG2/HBE1* could be partially distinct from the known association at the SCT variant. In our AA sample, 12 individuals carried two copies of rs1050828-T allele and at least one copy of rs1039215-G allele and had a mean HbA1c of 4.60 (0.41) *versus* 5.65 (0.39) for the 2,372 individuals that carried no risk allele at the two variants (**Supplementary Table 14**).

### HbA1c-lowering effects of *G6PD* variants in AA and HA

In sex-stratified analyses in AA, the *G6PD* rs76723693 variant was associated with a decrease in HbA1c of 0.98%-units (95% CI 0.51–1.44) per allele in hemizygous men and 0.46%-units (95% CI 0.26–0.66) in heterozygous women; whereas, the *G6PD* rs1050828 variant was associated with a decrease in HbA1c of 0.88%-units (95% CI 0.81–0.95) per allele in hemizygous men and 0.34%-units (95% CI 0.30–0.39) in heterozygous women (Table 2 **and** Figure 4). We detected heterogeneity in effect sizes between males and females for rs1050828 (P_het_=0.008) but not for rs76723693 (P_het_=0.19). In HA, the *G6PD* rs1050828 variant was associated with a decrease in HbA1c of 0.84%-units (95% CI 0.47–1.22) per allele in hemizygous men and 0.25%-units (95% CI 0.06–0.45) in heterozygous women (Table 3 **and** Figure 4). Upon adjusting on the sickle cell trait (SCT) variant, rs334, the association of these two SNVs did not attenuate (**Supplementary Table 12**).

**Figure 4:**
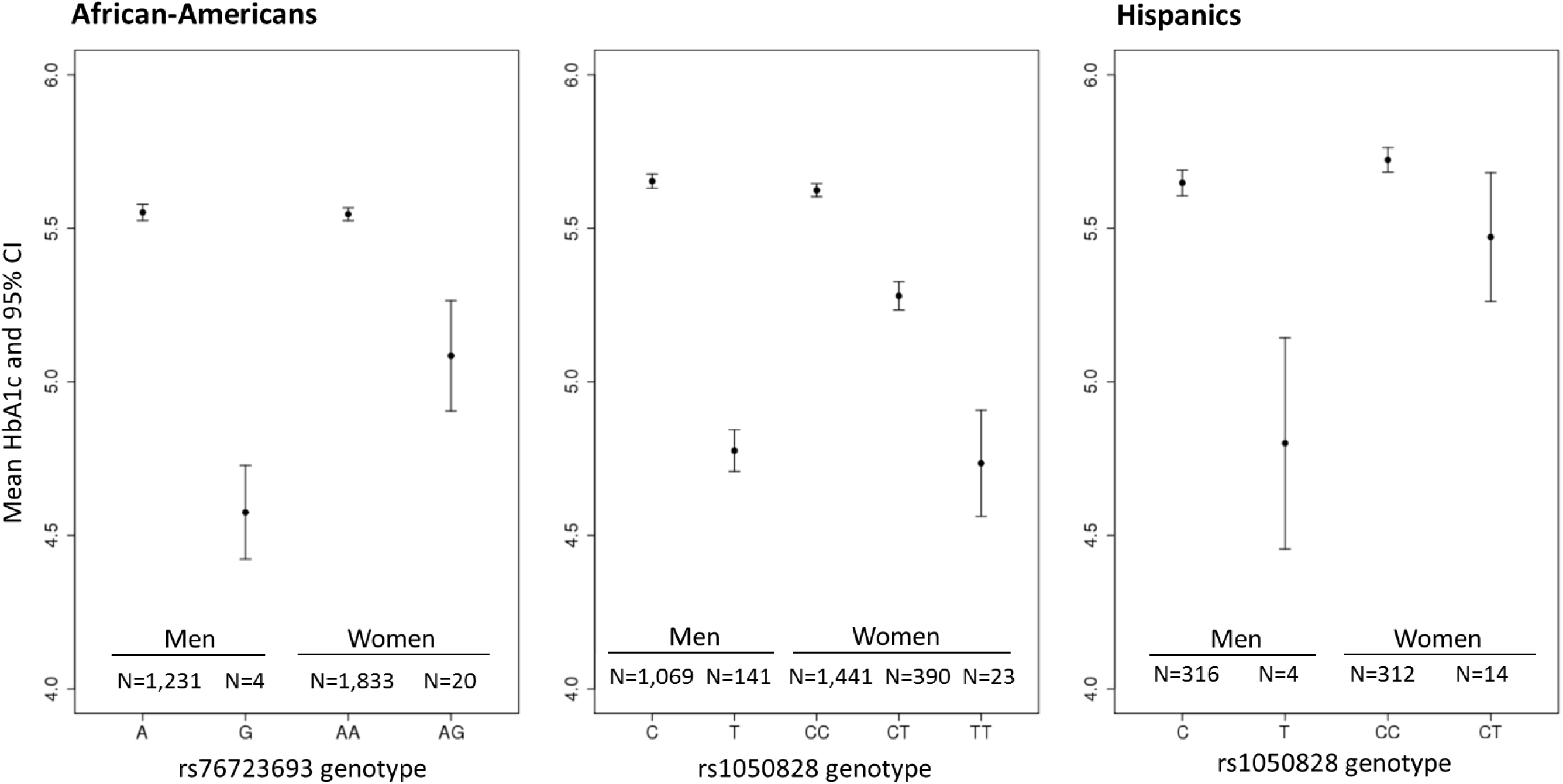
Mean HbA1c levels according to genotypes at both *G6PD* variants (rs76723693 and rs1050828) and stratified by sex. Plots are displayed separately in African-Americans (left and center) and in Hispanics (right). Due to the limited sample size of African-Americans women carrying two copies of the rare G allele at rs76723693, they are not represented on this plot. In AA, the *G6PD* rs76723693 variant was associated with a decrease in HbA1c of 0.98%-units (95% Confidence Interval (CI) 0.51–1.44) per allele in hemizygous men and 0.46%-units (95% CI 0.26–0.66) in heterozygous women; whereas, the *G6PD* rs1050828 variant was associated with a decrease in HbA1c of 0.88%-units (95% CI 0.81–0.95) per allele in hemizygous men and 0.34%-units (95% CI 0.30–0.39) in heterozygous women In HA, the *G6PD* rs1050828 variant was associated with a decrease in HbA1c of 0.84%-units (95% CI 0.47–1.22) per allele in hemizygous men and 0.25%-units (95% CI 0.06–0.45) in heterozygous women.

**Table 2:** Combined and sex-stratified conditional association analyses of HbA1c for the two distinct single nucleotide variants in the G6PD region in non-diabetic African-Americans in TOPMed cohorts

|  |  |  |  | Pooled analysis<br>(N=3,123) |  |  | Conditional analysis on<br>rs1050828 |  |  | Males (N=1,269) |  |  | Females (N=1,854) |  |  |  |
| --- | --- | --- | --- | --- | --- | --- | --- | --- | --- | --- | --- | --- | --- | --- | --- | --- |
| MarkerID<sup>c</sup> | rsID | Ref/Alt<br>Allele | MAF<sup>a</sup> | Beta | SE | *P value* | Beta | SE | *P value* | Beta | SE | *P value* | Beta | SE | *P value* | Heterogeneity<br>*P value*<sup>b</sup> |
| X:154533025 | rs76723693 | A/G | 0.005 | -0.44 | 0.07 | 2.2x10<sup>-9</sup> | -0.50 | 0.06 | 2.8x10<sup>-15</sup> | -0.52 | 0.12 | 6.8x10<sup>-6</sup> | -0.33 | 0.10 | 6.2x10<sup>-4</sup> | 0.19 |
| X:154536002 | rs1050828 | C/T | 0.12 | -0.41 | 0.01 | 8.4x10<sup>-205</sup> | - | - | - | -0.44 | 0.02 | 2.3x10<sup>-148</sup> | -0.37 | 0.02 | 2.7x10<sup>-69</sup> | 0.008 |

<sup>a</sup>MAF: Minor allele frequency
<sup>b</sup>Heterogeneity P value: test of difference in effect sizes between males and females, accounting for the correlation between males and females statistics
<sup>c</sup>MarkerID is defined as Chromosome:Position. Positions are indicated on Build 38.

**Table 3:** Analysis of HbA1c associations with rs1050828, X:154536002, alternate allele T, in non-diabetic African-Americans and Hispanics in TOPMed cohorts

|  | African-Americans |  |  |  |  |  | Hispanics |  |  |  |  |  |
| --- | --- | --- | --- | --- | --- | --- | --- | --- | --- | --- | --- | --- |
|  | N | MAF<sup>a</sup> | Beta | SE | *P value* | % of variance explained | N | MAF<sup>a</sup> | Beta | SE | *P value* | % of variance explained |
| All | 3,123 | 0.12 | -0.41 | 0.01 | 8.4E-205 | 0.23 | 650 | 0.02 | -0.37 | 0.07 | 8.3E-07 | 0.04 |
| Males | 1,269 | 0.12 | -0.44 | 0.02 | 2.3E-148 | 0.35 | 323 | 0.01 | -0.45 | 0.11 | 3.5E-05 | 0.05 |
| Females | 1,854 | 0.12 | -0.37 | 0.02 | 2.7E-69 | 0.14 | 327 | 0.02 | -0.28 | 0.1 | 4.5E-03 | 0.02 |

<sup>a</sup>MAF: Minor allele frequency

### Population reclassification of diabetes diagnosis by *G6PD* variants

To assess the potential public health impact of *G6PD* variants in AA, we designed a hypothetical scenario, using National Health and Nutrition Examination Survey (NHANES, see URL in the web resources section) 2015-2016, to estimate the number of African Americans and Hispanics whose diagnosis of diabetes would have been missed if screened with a single HbA1c measurement using the 6.5% diagnostic threshold and if the effects of the two *G6PD* variants rs1050828 and rs76723693 were not accounted for. We restricted the NHANES analytic sample to adults aged ≥ 18 years who self-identified as Non-Hispanic Black or Mexican American/Other Hispanic respectively. In non-Hispanic Blacks (N=1,227), an estimated 2.32% with HbA1c < 6.5%-units may be considered to have diabetes upon accounting for the effect size and observed allele frequency of the common *G6PD* variant, rs1050828. An additional 0.13% with HbA1c < 6.5%-units may be considered to have diabetes upon accounting for the effect of the rare *G6PD* variant, rs76723693. In Mexican American/Other Hispanic (N=1,768), an additional 0.26% with HbA1c < 6.5%-units may be considered to have diabetes due to the effect of rs1050828. According to the 2016 United States Census Bureau, approximately 30.14 and 39.33 million adults identified themselves as African American and Hispanic adults respectively, suggesting that 740,000 African American adults, of which 40,000 would be attributed to the rare variant, rs76723693, and 100,000 Hispanic adults with diabetes would remain undiagnosed when screened by a single HbA1c measurement if this genetic information was not taken into account (**Supplementary Table 15**).

### Replication of results in the UKBB

We selected a total of 5,964 African-ancestry non-diabetic UKBB participants for replication. In this sample, 2,461 participants were males, mean age (SD) was 51.1 (7.7), and mean HbA1c (SD) was 5.50 (0.44). The four variants of interest (rs334 (SCT), rs1039215 (HBG2), rs76723693 and rs1050828 (G6PD)) had good quality of imputation (info score > 0.50, **Supplementary Table 16**). We observed significant and consistent associations of the two *G6PD* variants with HbA1c (rs1050828-T, B=-0.36, P<2.2×10^-16^ and rs76723693-G, B=-0.21, P=0.007, **Supplementary Table 16**). The association of the rare *G6PD* rs76723693-G variant was even more significant when both *G6PD* variants were in the model (B=-0.27, P=6.7× 10^-5^). We did not detect significant associations of *HBG2* and *SCT* variants with HbA1c in the UKBB.

## Discussion

Through deep large-scale WGS association analysis in 10,338 individuals from four different ancestries and fine-mapping of HbA1c-associated loci, we identified common, low frequency and rare genetic variants that influence nonglycemic variation in HbA1c. We confirmed four known regions associated with HbA1c (the glycemic *GCK*, erythrocytic *HK1* and *G6PD*, and unclassified *FN3K/FN3KRP* regions) and discovered two new AA-specific low frequency or rare erythrocytic variants in *G6PD* (rs76723693) and *HBG2/HBE1* (rs1039215). The magnitude and direction of effect of the association of rs76723693 and rs1039215 with HbA1c was similar in the two largest TOPMed AA cohorts (JHS and MESA, **Supplementary Table 17**). The association of rs76723693 with HbA1c was replicated in UKBB African-ancestry participants. We detected significant association using gene-based test when aggregating rare missense variants in *G6PD*, indicating that others rare missense variants in *G6PD*, in addition to rs76723693, are associated with HbA1c. We showed that the association of the *HBG2/HBE1* variant with HbA1c was partially distinct from the known SCT rs334 variant in *HBB* [MIM:141900] suggesting that genetic variation at hemoglobin genes other than *HBB* may also influence HbA1c. Individuals carrying both the *G6PD* variant (rs1050828) and the *HBG2/HBE1* variant (rs1039215) had even lower HbA1c than those carrying only one of the two variants. Hemizygous males and homozygous females for the *G6PD* rs1050828-T allele who carried one or more copies of the *HBG2* rs1039215-G allele had a mean HbA1c that was 1.05%- unit lower than those carrying none of these alleles.

The novel HbA1c variant in AA, rs1039215 on chromosome 11, lies in an intron of the hemoglobin subunit gamma 2 gene *HBG2* and the hemoglobin subunit epsilon 1 gene *HBE1*. *HBG2,* in addition to *HBG1,* encodes the gamma chain of hemoglobin, which combines with 2 alpha chains to form fetal hemoglobin. Fetal hemoglobin is known to interfere with the measurement of HbA1c by some laboratory assays,^27–29^ and so persistence of fetal hemoglobin may be a mechanism through which rs1039215 influences HbA1c measurements. While a small degree of analytic interference with SCT has been reported for the Tosoh 2.2 and G7 assays used by JHS, MESA, and ARIC to measure HbA1c (see NGSP website URL in the web resources section),^30, 31^ no interference has been reported for the Bio-Rad Variant II Turbo assay used by UKB^32^ (**Supplementary Table 2**). Thus, assay interference may explain the lack of association of rs334 with HbA1c in UKBB. Alternatively, as our haplotype analysis indicated that the haplotype containing both rs334-A in *HBB* and rs1039215-G in *HBG2* was likely the main driver of this signal in the region, the true causal variant could be another variant within this haplotype that partially explains the reported association of rs334 with HbA1c.

Both variants identified in *G6PD* in AA are missense and pathogenic variants for G6PD deficiency (ClinVar), an X-linked genetic disorder characterized by a defect glucose-6-phosphate dehydrogenase enzymatic action, causing erythrocytes to break down prematurely. Thus, the mechanism through which these *G6PD* variants lower HbA1c is most likely through shortening of erythrocytic lifespan (**Supplementary Table 18**). Assuming random X-inactivation, we would expect the difference between hemizygous males and no rs1050828-T variant to be two times larger than the difference between heterozygous females and no rs1050828-T variant; however, in this study, we observed a slightly lower than expected effect in females. We posit that this heterogeneity of genetic effects by sex may be due to X-linked dosage compensation and not actual sex differences.^33^ While these *G6PD* variants lower HbA1c in AA, similar to sickle cell trait,^5^ we note that the lowering effect of these genetic variants on HbA1c do not explain the higher mean HbA1c in AA compared to European American with similar glycemia that has been reported in observational studies.^34, 35^

We found that rs1050828, the common *G6PD* variant in AA, was also associated with HbA1c in HA (sample composed of 51% Mexican, 14% Dominican, 14% Puerto Rican, 11% South American, and 10% others) suggesting that carriers of this variant are likely to be underdiagnosed if a single HbA1c measurement is used to screen two of the largest racial/ethnic minorities in the United States for diabetes. If *G6PD* variants were ignored, we estimate that approximately 840,000 African American and Hispanic adults with diabetes would remain undiagnosed. As Hispanic populations are ancestrally diverse including African Ancestry (~7% of HA in our study have > 50% AA ancestry and ~25% have > 15% AA ancestry), *G6PD* variants were expected to occur at some lower frequency in HA compared to AA. Given that there is no universal screening for G6PD deficiency in North America, asymptomatic carriers are unlikely to know their G6PD status.^7^ While misclassification of diabetes status at the population level was largely driven by the common G6PD variant, rs10105828, and not the rare rs76723693 variant (0.5% of the AA population), in this era of precision medicine there is a growing need for clinical practice to account for the impact of rare genetic variants with clinically meaningful effects on routinely performed diagnostic tests even if the proportion of carriers in the population is small. We also predict that other *G6PD* deficiency variants, particularly common in African, Mediterranean and Asian populations,^36^ will impact HbA1c’s utility as a diagnostic measure. Until genetic information is universally accessible to adjust diagnostic thresholds of imperfect biomarkers, our findings support current clinical practice guidelines by the American Diabetes Association which recommends the use of both FG and HbA1c in combination for diabetes diagnosis, especially given that the diagnostic level of HbA1c is conservative relative to FG.^37^

Strengths of this investigation include the analysis of extremely high-quality deep sequence data from a large, multi-ancestry sample including five studies and four ancestries to investigate the association of genetic variation across the full allelic spectrum with HbA1c, a diagnostic test that is central to the care of patients with diabetes. As previous GWAS on HbA1c had included a smaller sample of Hispanic ancestry, this study represents one of the first large-scale genetic discovery efforts for HbA1c that include multiple ethnic minorities in the United States. Our study demonstrates that both common and rare genetic variation differ across ancestries and informs the unique genetic architecture of HbA1c. We acknowledge limitations. Our hypothetical scenario using NHANES data to assess the population impact of *G6PD* genetics on diabetes screening does not take into account the full complement of clinical and genetic information contributing to nonglycemic variation in HbA1c, some of which may raise, and not lower, HbA1c reducing the risk of under-detection.^38^ For instance, structural variants (e.g., copy number variation) were not included in our analyses. Nevertheless, our findings highlight the potential for underdiagnosis in carriers of these variants, which disproportionately affect certain ethnic minorities, if HbA1c is used as the only diagnostic test for diabetes. Our sample sizes, and thus our power, are limited compared to GWAS, particularly for Hispanic and Asian populations. For instance, we did not detect an association signal for rs76723693 in HA because only one individual in our HA population carried a copy of the rare variant. Including larger samples of individuals of non-European ancestries will improve the characterization of rare variants with clinically meaningful effects on HbA1c. We acknowledge that the genome-wide threshold used in this paper may not represent the true number of independent genetic variants (SNVs as well as insertion-deletion variants) tested using our sequence data.

In this study we identified common and rare genetic variants that cause nonglycemic differences in HbA1c with important clinical and public health implications. Because HbA1c is commonly used in diabetes screening and management worldwide, disregarding the effects of these rare and common genetic variants that tend to occur in high diabetes-risk minority groups, could result in delayed diagnosis or under-detection of diabetes in carriers of these variants. To avoid such disparities in care, an assessment of these variants should be considered for incorporation into precision medicine approaches for diabetes diagnosis.

## Supplementary Data

Supplemental data contains cohort descriptions, 18 tables and 6 figures.

## Whole genome sequencing

Whole genome sequencing (WGS) for the Trans-Omics in Precision Medicine (TOPMed) program was supported by the National Heart, Lung and Blood Institute (NHLBI). WGS for “NHLBI TOPMed: Genetics of Cardiometabolic Health in the Amish (phs000956.v1.p1) was performed at the Broad Institute of MIT and Harvard (3R01HL121007-01S1). WGS for “NHLBI TOPMed: Trans-Omics for Precision Medicine Whole Genome Sequencing Project: ARIC (phs001211.v1.p1) was performed at the Broad Institute of MIT and Harvard and at the Baylor Human Genome Sequencing Center (3R01HL092577-06S1 (Broad, AFGen), HHSN268201500015C (Baylor, VTE), 3U54HG003273-12S2 (Baylor, VTE)). WGS for “NHLBI TOPMed: Whole Genome Sequencing and Related Phenotypes in the Framingham Heart Study (phs000974.v1.p1) was performed at the Broad Institute of MIT and Harvard (3R01HL092577-06S1 (AFGen)). WGS for “NHLBI TOPMed: The Jackson Heart Study (phs000964.v1.p1) was performed at the University of Washington Northwest Genomics Center (HHSN268201100037C). WGS for “NHLBI TOPMed: MESA and MESA Family AA-CAC (phs001416) was performed at the Broad Institute of MIT and Harvard (3U54HG003067-13S1 (MESA, TOPMed supplement to NHGRI), HHSN268201500014C (Broad, AA_CAC)). WGS for “NHLBI TOPMed: Cardiovascular Health Study (phs001368.v1.p1)” was performed at Baylor Human Genome Sequencing Center (HHSN268201500015C, VTE portion of CHS). WGS for “NHLBI TOPMed: GeneSTAR (Genetic Study of Atherosclerosis Risk) (phs001218.v1.p1)” was performed at Macrogen Corp and at the Broad Institute of MIT and Harvard (HHSN268201500014C (Broad, AA_CAC)). WGS for “NHLBI TOPMed: Women’s Health Initiative (WHI) (phs001237.v1.p1)” was performed at the Broad Institute of MIT and Harvard (HHSN268201500014C). WGS for “NHLBI TOPMed: San Antonio Family Heart Study (WGS) (phs001215.v1.p1)” was performed at Illumina Genomic Services (3R01HL113323-03S1). Centralized read mapping and genotype calling, along with variant quality metrics and filtering were provided by the TOPMed Informatics Research Center (3R01HL-117626-02S1). Phenotype harmonization, data management, sample-identity QC, and general study coordination, were provided by the TOPMed Data Coordinating Center (3R01HL-120393-02S1). We gratefully acknowledge the studies and participants who provided biological samples and data for TOPMed. The contributions of the investigators of the NHLBI TOPMed Consortium (https://www.nhlbiwgs.org/topmed-banner-authorship) are gratefully acknowledged.

## The Amish study

NIH grant R01 HL121007

## The Atherosclerosis Risk in Communities Study

Whole genome sequencing (WGS) for the Trans-Omics in Precision Medicine (TOPMed) program was supported by the National Heart, Lung and Blood Institute (NHLBI). WGS for “NHLBI TOPMed: Atherosclerosis Risk in Communities (ARIC)” (phs001211) was performed at the Baylor College of Medicine Human Genome Sequencing Center (HHSN268201500015C and 3U54HG003273-12S2) and the Broad Institute for MIT and Harvard (3R01HL092577-06S1). Centralized read mapping and genotype calling, along with variant quality metrics and filtering were provided by the TOPMed Informatics Research Center (3R01HL-117626-02S1). Phenotype harmonization, data management, sample-identity QC, and general study coordination, were provided by the TOPMed Data Coordinating Center (3R01HL-120393-02S1). We gratefully acknowledge the studies and participants who provided biological samples and data for TOPMed.

The Atherosclerosis Risk in Communities study has been funded in whole or in part with Federal funds from the National Heart, Lung, and Blood Institute, National Institutes of Health, Department of Health and Human Services (contract numbers HHSN268201700001I, HHSN268201700002I, HHSN268201700003I, HHSN268201700004I and HHSN268201700005I). The authors thank the staff and participants of the ARIC study for their important contributions.

## The Cardiovascular Health Study

This research was supported by contracts HHSN268201200036C, HHSN268200800007C, HHSN268201800001C, N01HC55222, N01HC85079, N01HC85080, N01HC85081, N01HC85082, N01HC85083, N01HC85086, and grants R01HL120393, U01HL080295 and U01HL130114 from the National Heart, Lung, and Blood Institute (NHLBI), with additional contribution from the National Institute of Neurological Disorders and Stroke (NINDS). Additional support was provided by R01AG023629 from the National Institute on Aging (NIA). A full list of principal CHS investigators and institutions can be found at CHS-NHLBI.org.

## The Framingham Heart Study

This research was conducted in part using data and resources from the Framingham Heart Study of the National Heart Lung and Blood Institute of the National Institutes of Health and Boston University School of Medicine. The Framingham Heart Study (FHS) acknowledges the support of contracts NO1-HC-25195 and HHSN268201500001I from the National Heart, Lung and Blood Institute, contract 75N92019D00031 and grant supplement R01 HL092577-06S1 for this research. This work was also supported in part by grant U01DK078616.

We also acknowledge the dedication of the FHS study participants without whom this research would not be possible.

## The Genetic Studies of Atherosclerosis Risk

GeneSTAR was supported by the National Institutes of Health/National Heart, Lung, and Blood Institute (U01 HL72518, HL087698, HL112064, HL11006, HL118356) and by a grant from the National Institutes of Health/National Center for Research Resources (M01-RR000052) to the Johns Hopkins General Clinical Research Center. We would like to thank the participants and families of GeneSTAR and our dedicated staff for all their sacrifices.

## The Jackson Heart Study

The Jackson Heart Study (JHS) is supported and conducted in collaboration with Jackson State University (HHSN268201800013I), Tougaloo College (HHSN268201800014I), the Mississippi State Department of Health (HHSN268201800015I/HHSN26800001) and the University of Mississippi Medical Center (HHSN268201800010I, HHSN268201800011I and HHSN268201800012I) contracts from the National Heart, Lung, and Blood Institute (NHLBI) and the National Institute for Minority Health and Health Disparities (NIMHD). The authors also wish to thank the staffs and participants of the JHS.

The views expressed in this manuscript are those of the authors and do not necessarily represent the views of the National Heart, Lung, and Blood Institute; the National Institutes of Health; or the U.S. Department of Health and Human Services.

## The Multi-Ethnic Study of Atherosclerosis

Whole genome sequencing (WGS) for the Trans-Omics in Precision Medicine (TOPMed) program was supported by the National Heart, Lung and Blood Institute (NHLBI). WGS for “NHLBI TOPMed: Multi-Ethnic Study of Atherosclerosis (MESA)” (phs001416.v1.p1) was performed at the Broad Institute of MIT and Harvard (3U54HG003067-13S1). Centralized read mapping and genotype calling, along with variant quality metrics and filtering were provided by the TOPMed Informatics Research Center (3R01HL-117626-02S1). Phenotype harmonization, data management, sample-identity QC, and general study coordination, were provided by the TOPMed Data Coordinating Center (3R01HL-120393-02S1). MESA and the MESA SHARe project are conducted and supported by the National Heart, Lung, and Blood Institute (NHLBI) in collaboration with MESA investigators. Support for MESA is provided by contracts HHSN268201500003I, N01-HC-95159, N01-HC-95160, N01-HC-95161, N01-HC-95162, N01-HC-95163, N01-HC-95164, N01-HC-95165, N01-HC-95166, N01-HC-95167, N01-HC-95168, N01-HC-95169, UL1-TR-000040, UL1-TR-001079, UL1-TR-001420. The provision of genotyping data was supported in part by the National Center for Advancing Translational Sciences, CTSI grant UL1TR001881, and the National Institute of Diabetes and Digestive and Kidney Disease Diabetes Research Center (DRC) grant DK063491 to the Southern California Diabetes Endocrinology Research Center.

## The San Antonio Family Study

Collection of the San Antonio Family Study data was supported in part by National Institutes of Health (NIH) grants R01 HL045522, DK047482, DK053889, MH078143, MH078111 and MH083824; and whole genome sequencing of SAFS subjects was supported by U01 DK085524 and R01 HL113323. This work was conducted in part in facilities constructed under the support of NIH grant C06 RR020547. We are very grateful to the participants of the San Antonio Family Study for their continued involvement in our research programs.

## The Women’s Health Initiative

The WHI program is funded by the National Heart, Lung, and Blood Institute, National Institutes of Health, U.S. Department of Health and Human Services through contracts HHSN268201600018C, HHSN268201600001C, HHSN268201600002C, HHSN268201600003C, and HHSN268201600004C. The authors thank the WHI investigators and staff for their dedication, and the study participants for making the program possible. A full listing of WHI investigators can be found at: http://www.whi.org/researchers/Documents%20%20Write%20a%20Paper/WHI%20Investigator%20Long%20List.pdf

## UKBB

This research has been conducted using the UK Biobank Resource under Application Number 42614.

## Web Resources

Analysis Commons: http://analysiscommons.com

EPACTS: https://genome.sph.umich.edu/wiki/EPACTS

HbA1c Assay Interferences information on the National Glycohemoglobin Standardization Program (NGSP) website: http://www.ngsp.org/interf.asp

National Health and Nutrition Examination Survey (NHANES): https://wwwn.cdc.gov/nchs/nhanes/continuousnhanes/default.aspx?BeginYear=2015

OASIS: https://omicsoasis.github.io

SmartPCA: http://www.hsph.harvard.edu/alkes-price/software

TOPMed Pipeline: https://github.com/UW-GAC/analysis_pipeline

TOPMed website: www.nhlbiwgs.org

Details regarding TOPMed laboratory methods, data processing and quality control: https://www.nhlbiwgs.org/methods

UK Biobank: https://www.ukbiobank.ac.uk/

## References

1. Mortensen, H.B., Christophersen, C. (1983). Glucosylation of human haemoglobin a in red blood cells studied in vitro. Kinetics of the formation and dissociation of haemoglobin A1c. Clin. Chim. Acta 134, 317–326.

2. Leong, A., Meigs, J.B. (2015). Type 2 Diabetes Prevention: Implications of Hemoglobin A1c Genetics. Rev. Diabet. Stud. 12, 351–362.

3. Wheeler, E., Leong, A., Liu, C.T., Hivert, M.F., Strawbridge, R.J., Podmore, C., Li, M., Yao, J., Sim, X., Hong, J. et al. (2017). Impact of common genetic determinants of Hemoglobin A1c on type 2 diabetes risk and diagnosis in ancestrally diverse populations: A transethnic genome-wide meta-analysis. PLoS Med. 14, e1002383.

4. International HapMap Consortium, Frazer, K.A., Ballinger, D.G., Cox, D.R., Hinds, D.A., Stuve, L.L., Gibbs, R.A., Belmont, J.W., Boudreau, A., Hardenbol, P. et al. (2007). A second generation human haplotype map of over 3.1 million SNPs. Nature 449, 851–861.

5. Lacy, M.E., Wellenius, G.A., Sumner, A.E., Correa, A., Carnethon, M.R., Liem, R.I., Wilson, J.G., Sacks, D.B., Jacobs, D.R., Jr, Carson, A.P., et al. (2017). Association of Sickle Cell Trait With Hemoglobin A1c in African Americans. JAMA 317, 507–515.

6. Strickland, S.W., Campbell, S.T., Little, R.R., Bruns, D.E., Bazydlo, L.A.L. (2018). Recognition of rare hemoglobin variants by hemoglobin A1c measurement procedures. Clin. Chim. Acta 476, 67–74.

7. Leong, A. (2007). Is there a need for neonatal screening of glucose-6-phosphate dehydrogenase deficiency in Canada? Mcgill J. Med. 10, 31–34.

8. Motulsky, A.G., Stamatoyannopoulos, G. (1966). Clinical implications of glucose-6-phosphate dehydrogenase deficiency. Ann. Intern. Med. 65, 1329–1334.

9. Paterson, A.D. (2017). HbA1c for type 2 diabetes diagnosis in Africans and African Americans: Personalized medicine NOW! PLoS Med. 14, e1002384.

10. Little, R.R., Rohlfing, C.L., Sacks, D.B., National Glycohemoglobin Standardization Program (NGSP) Steering Committee. (2011). Status of hemoglobin A1c measurement and goals for improvement: from chaos to order for improving diabetes care. Clin. Chem. 57, 205–214.

11. Regier, A.A., Farjoun, Y., Larson, D.E., Krasheninina, O., Kang, H.M., Howrigan, D.P., Chen, B.J., Kher, M., Banks, E., Ames, D.C. et al. (2018). Functional equivalence of genome sequencing analysis pipelines enables harmonized variant calling across human genetics projects. Nat. Commun. 9, 4038-018-06159-4.

12. Brody, J.A., Morrison, A.C., Bis, J.C., O’Connell, J.R., Brown, M.R., Huffman, J.E., Ames, D.C., Carroll, A., Conomos, M.P., Gabriel, S. et al. (2017). Analysis commons, a team approach to discovery in a big-data environment for genetic epidemiology. Nat. Genet. 49, 1560–1563.

13. Kanai, M., Tanaka, T., Okada, Y. (2016). Empirical estimation of genome-wide significance thresholds based on the 1000 Genomes Project data set. J. Hum. Genet. 61, 861–866.

14. Xu, C., Tachmazidou, I., Walter, K., Ciampi, A., Zeggini, E., Greenwood, C.M., UK10K Consortium. (2014). Estimating genome-wide significance for whole-genome sequencing studies. Genet. Epidemiol. 38, 281–290.

15. Astle, W.J., Elding, H., Jiang, T., Allen, D., Ruklisa, D., Mann, A.L., Mead, D., Bouman, H., Riveros-Mckay, F., Kostadima, M.A. et al. (2016). The Allelic Landscape of Human Blood Cell Trait Variation and Links to Common Complex Disease. Cell 167, 1415–1429.e19.

16. Hodonsky, C.J., Jain, D., Schick, U.M., Morrison, J.V., Brown, L., McHugh, C.P., Schurmann, C., Chen, D.D., Liu, Y.M., Auer, P.L. et al. (2017). Genome-wide association study of red blood cell traits in Hispanics/Latinos: The Hispanic Community Health Study/Study of Latinos. PLoS Genet. 13, e1006760.

17. Chen, Z., Tang, H., Qayyum, R., Schick, U.M., Nalls, M.A., Handsaker, R., Li, J., Lu, Y., Yanek, L.R., Keating, B. et al. (2013). Genome-wide association analysis of red blood cell traits in African Americans: the COGENT Network. Hum. Mol. Genet. 22, 2529–2538.

18. Lo, K.S., Wilson, J.G., Lange, L.A., Folsom, A.R., Galarneau, G., Ganesh, S.K., Grant, S.F., Keating, B.J., McCarroll, S.A., Mohler, E.R., 3rd et al. (2011). Genetic association analysis highlights new loci that modulate hematological trait variation in Caucasians and African Americans. Hum. Genet. 129, 307–317.

19. Chami, N., Chen, M.H., Slater, A.J., Eicher, J.D., Evangelou, E., Tajuddin, S.M., Love-Gregory, L., Kacprowski, T., Schick, U.M., Nomura, A. et al. (2016). Exome Genotyping Identifies Pleiotropic Variants Associated with Red Blood Cell Traits. Am. J. Hum. Genet. 99, 8–21.

20. Willer, C.J., Li, Y., Abecasis, G.R. (2010). METAL: fast and efficient meta-analysis of genomewide association scans. Bioinformatics 26, 2190–2191.

21. Liu, X., White, S., Peng, B., Johnson, A.D., Brody, J.A., Li, A.H., Huang, Z., Carroll, A., Wei, P., Gibbs, R. et al. (2016). WGSA: an annotation pipeline for human genome sequencing studies. J. Med. Genet. 53, 111–112.

22. Bycroft, C., Freeman, C., Petkova, D., Band, G., Elliott, L.T., Sharp, K., Motyer, A., Vukcevic, D., Delaneau, O., O’Connell, J. et al. (2018). The UK Biobank resource with deep phenotyping and genomic data. Nature 562, 203–209.

23. Patterson, N., Price, A.L., Reich, D. (2006). Population structure and eigenanalysis. PLoS Genet. 2, e190.

24. Price, A.L., Patterson, N.J., Plenge, R.M., Weinblatt, M.E., Shadick, N.A., Reich, D. (2006). Principal components analysis corrects for stratification in genome-wide association studies. Nat. Genet. 38, 904–909.

25. Purcell, S., Neale, B., Todd-Brown, K., Thomas, L., Ferreira, M.A., Bender, D., Maller, J., Sklar, P., de Bakker, P.I., Daly, M.J. et al. (2007). PLINK: a tool set for whole-genome association and population-based linkage analyses. Am. J. Hum. Genet. 81, 559–575.

26. Shriner, D., Rotimi, C.N. (2018). Whole-Genome-Sequence-Based Haplotypes Reveal Single Origin of the Sickle Allele during the Holocene Wet Phase. Am. J. Hum. Genet. 102, 547–556.

27. Little, R.R., Roberts, W.L. (2009). A review of variant hemoglobins interfering with hemoglobin A1c measurement. J. Diabetes Sci. Technol. 3, 446–451.

28. Rohlfing, C.L., Connolly, S.M., England, J.D., Hanson, S.E., Moellering, C.M., Bachelder, J.R., Little, R.R. (2008). The effect of elevated fetal hemoglobin on hemoglobin A1c results: five common hemoglobin A1c methods compared with the IFCC reference method. Am. J. Clin. Pathol. 129, 811–814.

29. Sarnowski, C., Hivert, M.F. (2018). Impact of Genetic Determinants of HbA1c on Type 2 Diabetes Risk and Diagnosis. Curr. Diab Rep. 18, 52-018-1022-4.

30. Roberts, W.L., Safar-Pour, S., De, B.K., Rohlfing, C.L., Weykamp, C.W., Little, R.R. (2005). Effects of hemoglobin C and S traits on glycohemoglobin measurements by eleven methods. Clin. Chem. 51, 776–778.

31. Rohlfing, C., Hanson, S., Little, R.R. (2017). Measurement of Hemoglobin A1c in Patients With Sickle Cell Trait. JAMA 317, 2237.

32. Mongia, S.K., Little, R.R., Rohlfing, C.L., Hanson, S., Roberts, R.F., Owen, W.E., D’Costa, M.A., Reyes, C.A., Luzzi, V.I., Roberts, W.L. (2008). Effects of hemoglobin C and S traits on the results of 14 commercial glycated hemoglobin assays. Am. J. Clin. Pathol. 130, 136–140.

33. Sidorenko, J., Kassam, I., Kemper, K., Zeng, J., Lloyd-Jones, L., Montgomery, G.W., Gibson, G., Metspalu, A., Esko, T., Yang, J. et al. (2018). The effect of X-linked dosage compensation on complex trait variation. bioRxiv.

34. Bergenstal, R.M., Gal, R.L., Connor, C.G., Gubitosi-Klug, R., Kruger, D., Olson, B.A., Willi, S.M., Aleppo, G., Weinstock, R.S., Wood, J. et al. (2017). Racial Differences in the Relationship of Glucose Concentrations and Hemoglobin A1c Levels. Ann. Intern. Med. 167, 95–102.

35. Ziemer, D.C., Kolm, P., Weintraub, W.S., Vaccarino, V., Rhee, M.K., Twombly, J.G., Narayan, K.M., Koch, D.D., Phillips, L.S. (2010). Glucose-independent, black-white differences in hemoglobin A1c levels: a cross-sectional analysis of 2 studies. Ann. Intern. Med. 152, 770–777.

36. Gomez-Manzo, S., Marcial-Quino, J., Vanoye-Carlo, A., Serrano-Posada, H., Ortega-Cuellar, D., Gonzalez-Valdez, A., Castillo-Rodriguez, R.A., Hernandez-Ochoa, B., Sierra-Palacios, E., Rodriguez-Bustamante, E. et al. (2016). Glucose-6-Phosphate Dehydrogenase: Update and Analysis of New Mutations around the World. Int. J. Mol. Sci. 17, 10.3390/ijms17122069.

37. American Diabetes Association. (2019). 2. Classification and Diagnosis of Diabetes: Standards of Medical Care in Diabetes-2019. Diabetes Care 42, S13–S28.

38. Jun, G., Sedlazeck, F.J., Chen, H., Yu, B., Qi, Q., Krasheninina, O., Carroll, A., Liu, X., Mansfield, A., Zarate, S. et al. (2018). PgmNr 3186/W: Identification of novel structural variations affecting common and complex disease risks with >16,000 whole genome sequences from ARIC and HCHS/SOL.

