## Supplementary Text and Figures for "Impact of rare and common genetic variants on diabetes diagnosis by hemoglobin A1c in multi-ancestry cohorts: The Trans-Omics for Precision Medicine Program"

Sarnowski C, Leong A et al

**Supplementary Information**

**Cohort Description**

1. **Cohorts included in the HbA1c analyses (and RBC analyses)**

**The Amish study**

The Amish Complex Disease Research Program (http://www.medschool.umaryland.edu/endocrinology/Amish-Research-Program/) includes a set of large community-based studies focused largely on cardiometabolic health carried out in the Old Order Amish (OOA) community of Lancaster County, Pennsylvania. Over 7,000 Amish have been recruited to date. This Amish community is a founder population who immigrated to Pennsylvania from Western Europe in the early 1700's, later expanding into other regions of the U.S. The Amish cohort participating in the TOPMed Consortium comprises 1,120 subjects ≥ 18 years of age from large multigenerational families who were recruited for specific protocols between 2001 and 2006. Subjects have been extensively phenotyped for a range of cardiometabolic traits, including anthropometry, lipids, blood pressure, glucose and related measures, vascular imaging, and a range of other phenotypes. DNA samples have been collected and serum and plasma samples biobanked. The TOPMed Program has provided WGS data to complement GWAS array data already collected in >5,000 Amish study participants. Due to their ancestral history, the OOA are enriched for rare exonic variants that arose in the population from a single founder (or small number of founders) and propagated through genetic drift. Many of these variants have large effect sizes, and identifying them can lead to new biological insights about health and disease. A major goal of the TOPMed WGS sequencing efforts is to identify functional variants that underlie some of the large effect associations observed in this unique population.

Complete Blood Counts analyses were performed within 24 h for all individuals and consisted of RBC, HB, HCT, MCV, MCH, MCHC, RDW and WBC. RBC, HB, and MCV were directly measured, whereas HCT, MCH, and MCHC were mathematically derived from directly measured erythrocyte traits (MCV × RBC, HB/RBC, and MCH/MCV, respectively).[1]

**The Atherosclerosis Risk in Communities Study**

The ARIC study is a population-based prospective cohort study of cardiovascular disease sponsored by the National Heart, Lung, and Blood Institute (NHLBI). ARIC included 15,792 individuals, predominantly European American and African American, aged 45-64 years at baseline (1987-89) and chosen by probability sampling from four US communities. Cohort members completed three additional triennial follow-up examinations, a fifth exam in 2011-2013, and a sixth exam in 2016-2017. The ARIC study has been described in detail previously.[2]

Complete Blood Counts was measured using automated hematology analyzers: Coulter S + IV (calibration S - Cal, Beckman Coulter, Inc, Fullerton, CA) at 2 sites, Coulter S + III and Coulter S + IV (calibration S-Cal) at 1 site, and Technicon H-6000 (calibration Fisher, Technicon Corporation, Tarrytown, NY) at 1 site.

**The Framingham Heart Study**

The Framingham Heart Study (FHS) is a single-site, community-based, prospective cohort study that was initiated in 1948 to investigate risk factors for cardiovascular disease. The population of Framingham was almost entirely white in 1948. The FHS comprises three generations of participants: the original cohort followed since 1948 (Original or Gen1);[3] their offspring and spouses of the offspring, followed since 1971 (Offspring or Gen2);[4] and children from the largest offspring families enrolled in 2002 (Gen3).[5] The Original cohort enrolled 5,209 men and women who comprised two-thirds of the adult population then residing in Framingham, MA, USA. Survivors continue to receive biennial examinations. The Offspring cohort comprises 5,124 persons (including 3,514 biological offspring) who have been examined approximately once every 4 to 8 years and have completed 9 examinations. The first examination of the Gen3 was completed in July 2005 and included 4,095 participants, a third examination is currently ongoing. All cohorts continue under active surveillance for cardiovascular events. All participants provided written informed consent. This study was approved by the Institutional Review Board of the Boston University Medical Center.

The determination of hematological phenotypes in the FHS has been detailed previously.[6, 7] Hematology testing was performed on a Baker 9000 Hematology Analyzer (Baker Instruments Corp.) for the Original cohort and on a Beckman Coulter HmX Hematology Analyzer (Beckman Coulter, Inc.) for the Offspring and Gen3 cohorts.

**The Jackson Heart Study**

The Jackson Heart Study (JHS) was designed to study the reasons for the greater prevalence of cardiovascular disease among African Americans and to find new approaches for reducing this health disparity. JHS recruited 5,306 AA participants from urban and rural areas of the three counties (Hinds, Madison and Rankin) that comprise the Jackson, Mississippi metropolitan area from 2000-2004. Recruitment was limited to non-institutionalized adult African Americans. Participants were recruited in four ways: (1) randomly sampling households from a commercial listing; (2) a structured volunteer sample designed to mirror the eligible population; (3) current enrollment in the Atherosclerosis Risk in Communities (ARIC) study; and (4) a nested family cohort. Unrelated participants were between 35 and 84 years old, while members of the family cohort were $\geq$ 21 years old at baseline. A range of measures, including health behaviors, medication use, anthropometry, blood pressure, assessments of kidney function and diabetes, and CVD biomarkers, were assessed at the baseline JHS visit. The JHS has been described in detail previously.[8, 9] Blood cell count metrics were assessed using standard methods (Beckman Coulter automated hematology analyzer), as described previously.[10]

**The Multi-Ethnic Study of Atherosclerosis**

The Multi-Ethnic Study of Atherosclerosis (MESA) is a study of the characteristics of subclinical cardiovascular disease and the risk factors that predict progression to clinically overt cardiovascular disease or progression of the subclinical disease. MESA consisted of a diverse, population-based sample of an initial 6,814 asymptomatic men and women aged 45-84. 38 percent of the recruited participants were white, 28 percent African American, 22 percent Hispanic, and 12 percent Asian, predominantly of Chinese descent. Participants were recruited from six field centers across the United States: Wake Forest University, Columbia University, Johns Hopkins University, University of Minnesota, Northwestern University and University of California - Los Angeles. Each participant received an extensive physical exam and determination of coronary calcification, ventricular mass and function, flow-mediated endothelial vasodilation, carotid intimal-medial wall thickness and presence of echogenic lucencies in the carotid artery, lower extremity vascular insufficiency, arterial wave forms, electrocardiographic (ECG) measures, standard coronary risk factors, sociodemographic factors, lifestyle factors, and psychosocial factors. Selected repetition of subclinical disease measures and risk factors at follow-up visits allowed study of the progression of disease. Participants are being followed for identification and characterization of cardiovascular disease events, including acute myocardial infarction and other forms of coronary heart disease (CHD), stroke, and congestive heart failure; for cardiovascular disease interventions; and for mortality. The first examination took place over two years, from July 2000 - July 2002. It was followed by four examination periods that were 17-20 months in length. Participants have been contacted every 9 to 12 months throughout the study to assess clinical morbidity and mortality.

**MESA Family**

In the MESA Family Study, the goal is to locate and identify genes contributing to the genetic risk for cardiovascular disease (CVD), by looking at the early changes of atherosclerosis within families (mainly siblings). 2128 individuals from 594 families, yielding 3,026 sib pairs divided between African Americans and Hispanic-Americans, were recruited by utilizing the existing framework of MESA. MESA Family studied siblings of index subjects from the MESA study and from new sibpair families (with the same demographic characteristics) and is determining the extent of genetic contribution to the variation in coronary calcium (obtained via CT Scan) and carotid artery wall thickness (B-mode ultrasound) in the two largest non-majority U.S. populations. The MESA Family cohort was recruited from the six MESA Field Centers. MESA Family participants underwent the same examination as MESA participants during May 2004 - May 2007. The MESA Study has been described in detail previously.[11] Complete Blood Counts was performed at LabCorp using automated cell counter with mixed technologies.

2. **Cohorts included in the RBC analyses only**

**The Cardiovascular Health Study**

CHS is a population-based cohort study of risk factors for coronary heart disease and stroke in adults ≥65 years conducted across four field centers.[12] The original predominantly European ancestry cohort of 5,201 persons was recruited in 1989-1990 from random samples of the Medicare eligibility lists; subsequently, an additional predominantly African-American cohort of 687 persons was enrolled for a total sample of 5,888. CHS participants selected for inclusion in the TOPMed sequencing program included African-Americans participants, cases of idiopathic venous thromboembolism, myocardial infarction, coronary heart disease or stroke and a random sample of “healthy elderly”. Blood samples were drawn from all participants at their baseline examination and DNA was subsequently extracted from available samples. Details of measurement of hematologic traits have been previously described.[13]

CHS was approved by institutional review committees at each field center and individuals in the present analysis had available DNA and gave informed consent including consent to use of genetic information for the study of cardiovascular disease.

**The Genetic Studies of Atherosclerosis Risk**

GeneSTAR: The Genetic Study of Atherosclerosis Risk (GeneSTAR) is a longitudinal family-based study designed to explore environmental, phenotypic, and genetic causes of premature cardiovascular disease. European- and African-American participants were recruited from European- and African-American families (n=891) identified from probands hospitalized for a coronary disease event prior to 60 years of age in any of 10 Baltimore, Maryland area hospitals between 1983 and 2006. Apparently healthy siblings of the probands, offspring of the siblings and probands, and the co-parents of the offspring were screened for traditional coronary disease and stroke risk factors between 2003 and 2006 as part of a platelet function study of two weeks of 81 mg/day of aspirin.[14, 15]. All measures described here were obtained prior to the commencement of aspirin. Exclusion criteria included: 1) any coronary heart disease or vascular thrombotic event, 2) any bleeding disorder or hemorrhagic event, 3) current use of any anticoagulants or anti-platelet agents, 4) current use of chronic or acute nonsteroidal anti-inflammatory agents that could not be discontinued, 5) recent active gastrointestinal disorder, 6) current pharmacotherapy for a gastrointestinal disorder, 7) pregnancy or risk of pregnancy during the trial, 8) recent menorrhagia, 9) known aspirin intolerance or allergic side effects, 10) serious medical disorders, (eg, autoimmune diseases, cancer or HIV-AIDS), 11) current chronic or acute use of glucocorticosteroid therapy or any drug that may interfere with the measured outcomes, 12) serious psychiatric disorders, and, 13) inability to independently make a decision to participate. Of 3003 aspirin study participants, 1786 were selected for TOPMed based on 1) complete platelet function phenotyping and 2) largest family size. Complete Cell Counts were obtained using an automated cell counter (ACT-Diff, Beckman Coulter).

**The San Antonio Family Study**

The San Antonio Family Study results from the amalgamation of two San Antonio-based genetic studies. The first is the longitudinal San Antonio Family Heart Study (SAFHS) which began in 1991 and was designed to primarily investigate the genetics of cardiovascular disease and its risk factors in Mexican Americans. The SAFHS included 1,431 individuals in 42 extended families at baseline.[16] With some additional recruiting, it has now been expanded to 1,662 individuals in 47 families. Ascertainment occurred by way of the random selection of an adult Mexican American proband, without regard to presence or absence of disease and almost exclusively from Mexican American census tracts in San Antonio. The second component study is the San Antonio Family Gall Bladder Study (Dr. Duggirala, PI) which included 907 individuals from 39 families ascertained similarly to the SAFHS but with the requirement that the original proband also be diabetic.[17] This is a very weak form of ascertainment in Mexican Americans, where lifetime prevalence of diabetes approaches 30%. In fact, 20 years after the initiation of the SAFS, the prevalence for major diseases such as heart disease, diabetes, and obesity are not significantly different between these two component studies. Finally, we have expanded these pedigrees in recent years by examining a set of 498 children than are part of these families. This expansion was part of Dr. Duggirala's San Antonio Family Assessment of Metabolic Risk Factors in Youth (SAFARI) study. Additionally, we have seen 112 of these children subsequently as adults. Combined, these studies have 3,099 individuals primarily from 73 families. Our study is a mixed longitudinal design. Subjects have been seen between 1 and 4 times with an average of 1.95 examinations. For this study, samples and data analyzed were from the San Antonio Family Heart Study component. Blood cell counts were obtained within 2hrs of blood collection from an EDTA tube using a Beckman Coulter AcT diff2 Hematology Analyzer.

**The Women's Health Initiative**

The Women's Health Initiative (WHI) is a long-term, prospective, multi-center cohort study investigating post-menopausal women's health in the US.[18] WHI was funded by the National Institutes of Health and the National Heart, Lung, and Blood Institute to study strategies to prevent heart disease, breast cancer, colon cancer, and osteoporotic fractures in women 50-79 years of age. WHI involves 161,808 women recruited between 1993 and 1998 at 40 centers across the US. The study consists of two parts: the WHI Clinical Trial which was a randomized clinical trial of hormone therapy, dietary modification, and calcium/Vitamin D supplementation, and the WHI Observational Study, which focused on many of the inequities in women's health research and provided practical information about incidence, risk factors, and interventions related to heart disease, cancer, and osteoporotic fractures. Fasting blood samples were drawn and analyzed for RBC traits at designated clinical laboratories using an automated electronic cell counter at the baseline examination.[19] Participants with HCT>80, HGB>30, RBC=0 or HCT/HGB>7 were excluded. In total, 9912, 9909 and 1285 participants were included for HGB, HCT and RDW analyses, respectively, and 1286 participants were included for MCH, MCHC, MCV and RBC counts analyses.

### References

1. Hinckley, J.D., Abbott, D., Burns, T.L., Heiman, M., Shapiro, A.D., Wang, K., Di Paola, J. (2013). Quantitative trait locus linkage analysis in a large Amish pedigree identifies novel candidate loci for erythrocyte traits. Mol. Genet. Genomic Med. *1*, 131-141.

2. Anonymous (1989). The Atherosclerosis Risk in Communities (ARIC) Study: design and objectives. The ARIC investigators. Am. J. Epidemiol. *129*, 687-702.

3. Dawber, T.R., Kannel, W.B. (1966). The Framingham study. An epidemiological approach to coronary heart disease. Circulation *34*, 553-555.

4. Feinleib, M., Kannel, W.B., Garrison, R.J., McNamara, P.M., Castelli, W.P. (1975). The Framingham Offspring Study. Design and preliminary data. Prev. Med. *4*, 518-525.

5. Splansky, G.L., Corey, D., Yang, Q., Atwood, L.D., Cupples, L.A., Benjamin, E.J., D'Agostino RB, S., Fox, C.S., Larson, M.G., Murabito, J.M. et al. (2007). The Third Generation Cohort of the National Heart, Lung, and Blood Institute's Framingham Heart Study: design, recruitment, and initial examination. Am. J. Epidemiol. *165*, 1328-1335.

6. Yang, Q., Kathiresan, S., Lin, J.P., Tofler, G.H., O'Donnell, C.J. (2007). Genome-wide association and linkage analyses of hemostatic factors and hematological phenotypes in the Framingham Heart Study. BMC Med. Genet. *8 Suppl 1*, S12-2350-8-S1-S12.

7. Sloan, A., Gona, P., Johnson, A.D. (2015). Cardiovascular correlates of platelet count and volume in the Framingham Heart Study. Ann. Epidemiol. *25*, 492-498.

8. Taylor, H.A.,Jr, Wilson, J.G., Jones, D.W., Sarpong, D.F., Srinivasan, A., Garrison, R.J., Nelson, C., Wyatt, S.B. (2005). Toward resolution of cardiovascular health disparities in African Americans: design and methods of the Jackson Heart Study. Ethn. Dis. *15*, S6-4-17.

9. Wilson, J.G., Rotimi, C.N., Ekunwe, L., Royal, C.D., Crump, M.E., Wyatt, S.B., Steffes, M.W., Adeyemo, A., Zhou, J., Taylor, H.A.,Jr et al. (2005). Study design for genetic analysis in the Jackson Heart Study. Ethn. Dis. *15*, S6-30-37.

10. Carpenter, M.A., Crow, R., Steffes, M., Rock, W., Heilbraun, J., Evans, G., Skelton, T., Jensen, R., Sarpong, D. (2004). Laboratory, reading center, and coordinating center data management methods in the Jackson Heart Study. Am. J. Med. Sci. *328*, 131-144.

11. Bild, D.E., Bluemke, D.A., Burke, G.L., Detrano, R., Diez Roux, A.V., Folsom, A.R., Greenland, P., Jacob, D.R.,Jr, Kronmal, R., Liu, K. et al. (2002). Multi-Ethnic Study of Atherosclerosis: objectives and design. Am. J. Epidemiol. *156*, 871-881.

12. Fried, L.P., Borhani, N.O., Enright, P., Furberg, C.D., Gardin, J.M., Kronmal, R.A., Kuller, L.H., Manolio, T.A., Mittelmark, M.B., Newman, A. (1991). The Cardiovascular Health Study: design and rationale. Ann. Epidemiol. *1*, 263-276.

13. Ganesh, S.K., Zakai, N.A., van Rooij, F.J., Soranzo, N., Smith, A.V., Nalls, M.A., Chen, M.H., Kottgen, A., Glazer, N.L., Dehghan, A. et al. (2009). Multiple loci influence erythrocyte phenotypes in the CHARGE Consortium. Nat. Genet. *41*, 1191-1198.

14. Bray, P.F., Mathias, R.A., Faraday, N., Yanek, L.R., Fallin, M.D., Herrera-Galeano, J.E., Wilson, A.F., Becker, L.C., Becker, D.M. (2007). Heritability of platelet function in families with premature coronary artery disease. J. Thromb. Haemost. *5*, 1617-1623.

15. Faraday, N., Yanek, L.R., Mathias, R., Herrera-Galeano, J.E., Vaidya, D., Moy, T.F., Fallin, M.D., Wilson, A.F., Bray, P.F., Becker, L.C. et al. (2007). Heritability of platelet responsiveness to aspirin in activation pathways directly and indirectly related to cyclooxygenase-1. Circulation *115*, 2490-2496.

16. Mitchell, B.D., Kammerer, C.M., Blangero, J., Mahaney, M.C., Rainwater, D.L., Dyke, B., Hixson, J.E., Henkel, R.D., Sharp, R.M., Comuzzie, A.G. et al. (1996). Genetic and environmental contributions to cardiovascular risk factors in Mexican Americans. The San Antonio Family Heart Study. Circulation *94*, 2159-2170.

17. Duggirala, R., Blangero, J., Almasy, L., Dyer, T.D., Williams, K.L., Leach, R.J., O'Connell, P., Stern, M.P. (1999). Linkage of type 2 diabetes mellitus and of age at onset to a genetic location on chromosome 10q in Mexican Americans. Am. J. Hum. Genet. *64*, 1127-1140.

18. Anonymous (1998). Design of the Women's Health Initiative clinical trial and observational study. The Women's Health Initiative Study Group. Control. Clin. Trials *19*, 61-109.

19. Chen, Z., Tang, H., Qayyum, R., Schick, U.M., Nalls, M.A., Handsaker, R., Li, J., Lu, Y., Yanek, L.R., Keating, B. et al. (2013). Genome-wide association analysis of red blood cell traits in African Americans: the COGENT Network. Hum. Mol. Genet. *22*, 2529-2538.

**Supplementary Figure 1:** Quantile-Quantile (QQ)-plots by ancestry of the association analysis of HbA1c in non-diabetic individuals in TOPMed cohorts stratified on minor allele frequency

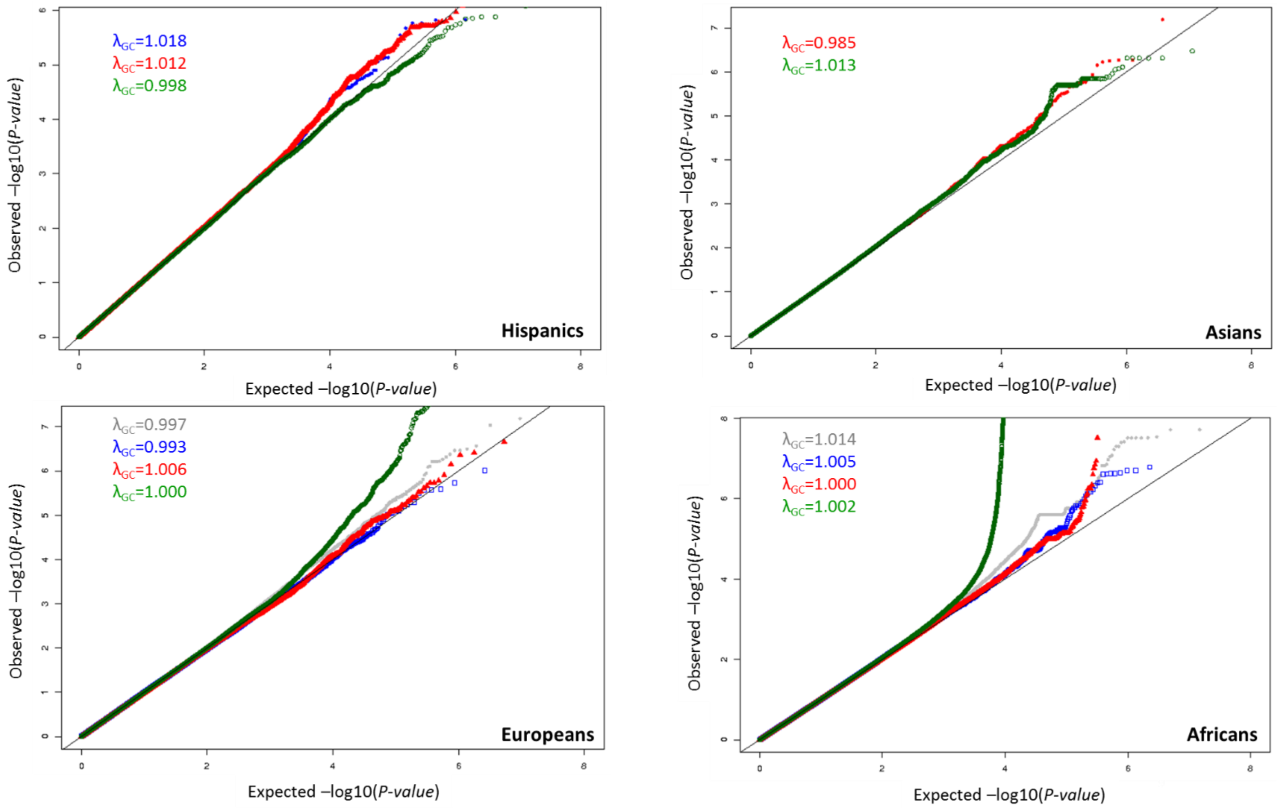


The dots represent the distribution of observed ordered –log10(*P-values*) against the theoretical model distribution of expected ordered –log10(*P-values*). The solid black line represents the theoretical model distribution of expected –log10(P-values) under the null distribution. The genomic inflation factor ($\lambda_{GC}$) is defined as the ratio of the median of the empirically observed distribution of the test statistic to the expected median, thus quantifying the extent of the inflation and the excess false positive rate. Association results were stratified by minor allele frequency (MAF) of single nucleotide variants: $MAF \geq 5\%$ are indicated in green; $5\% > MAF \geq 1\%$ are indicated in red; $1\% > MAF \geq 0.5\%$ are indicated in blue; $0.5\% > MAF \geq 0.1\%$ are indicated in grey.

**Supplementary Figure 2:** Manhattan-plots by ancestry of the association analysis of HbA1c in non-diabetic individuals in TOPMed cohorts.

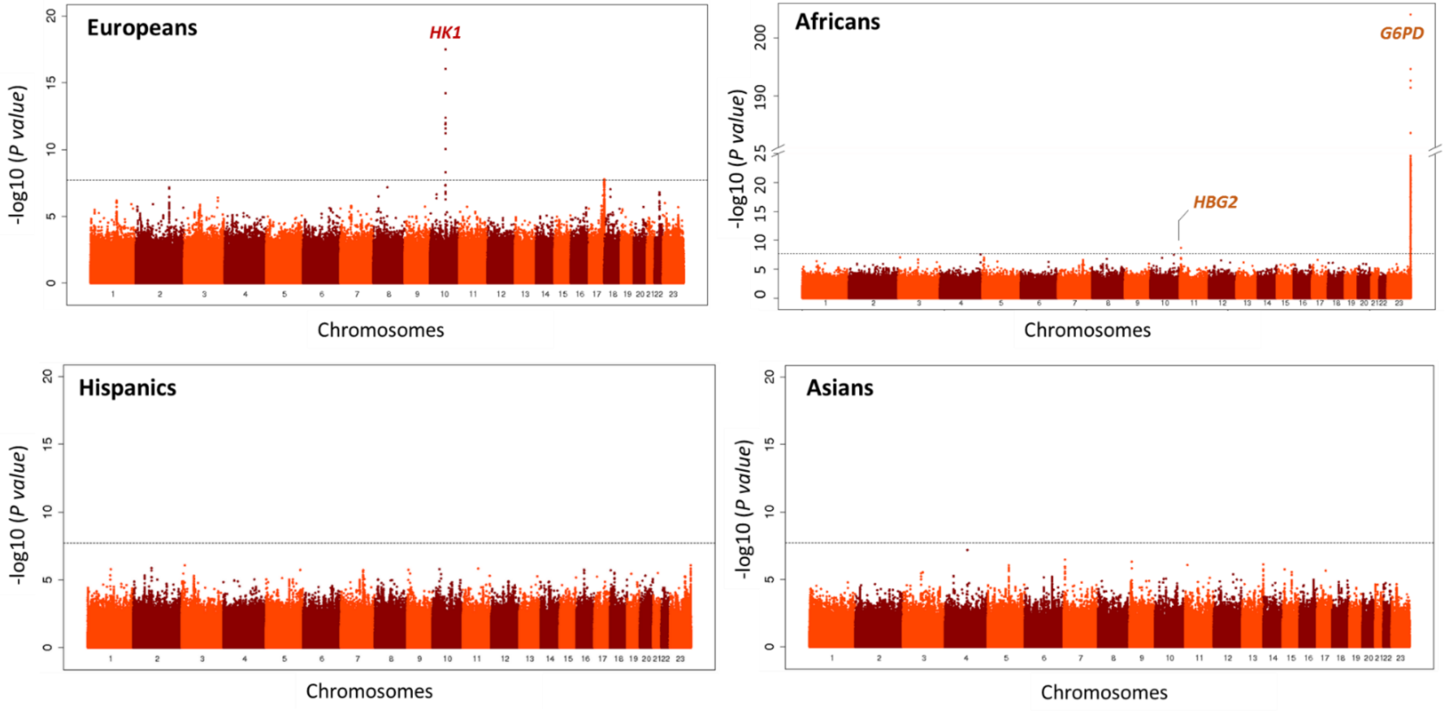


The –log10(P-value) for each single nucleotide variant on the y-axis is plotted against the build 38 genomic position on the x-axis (chromosomal coordinate). The dashed horizontal line indicates the genome-wide significance threshold of $P = 2\times10^{-8}$. The y-axis was truncated for ease of interpretation.

**Supplementary Figure 3**: Regional HbA1c association plot in the *HBG2/HBE1* region (+/- 750kb around *HBG2*).

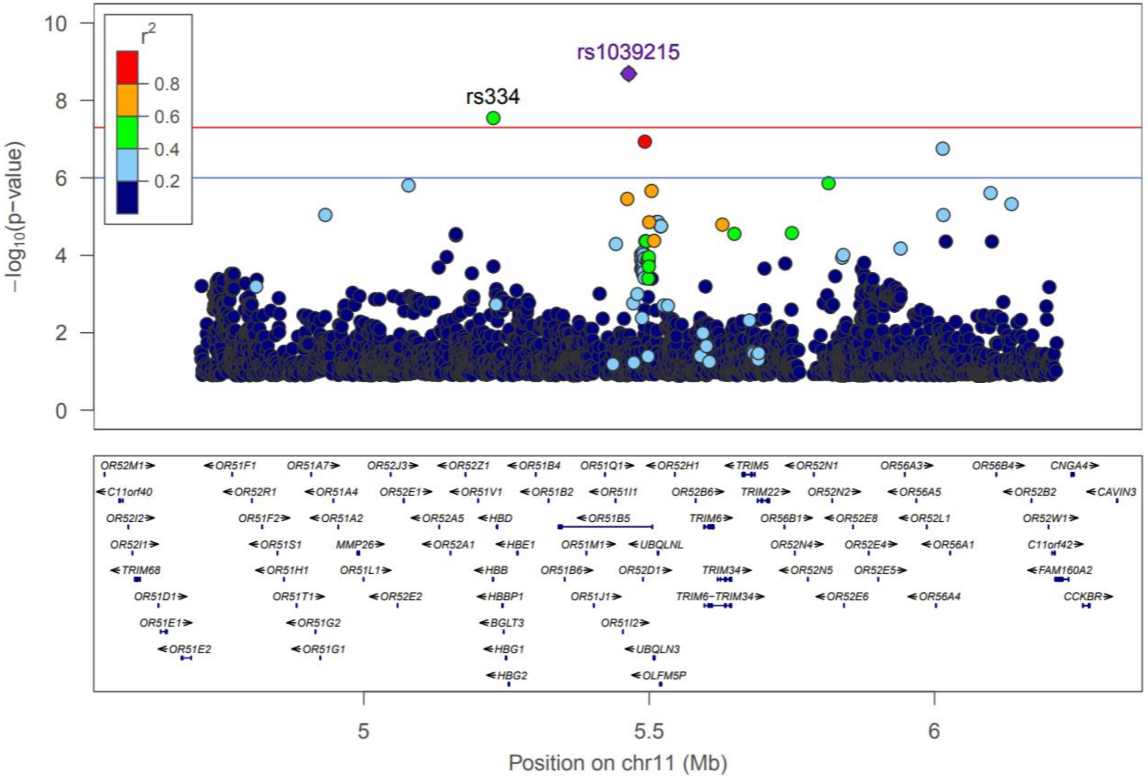


Single nucleotide variants are plotted with their P-values (-log10 values, left y-axis) as a function of build 38 genomic position on chromosome 11 (x-axis). Estimated recombination rates (right y-axis) are plotted to reflect the local linkage disequilibrium (LD) structure around the top associated single nucleotide variant rs1039215 (purple diamond) and correlated proxies (according to a blue to red scale from r2=0 to 1). LD was calculated in non-diabetic African ancestry TOPMed cohorts.

**Supplementary Figure 4**: Regional HbA1c association plot in the 10q25 region.

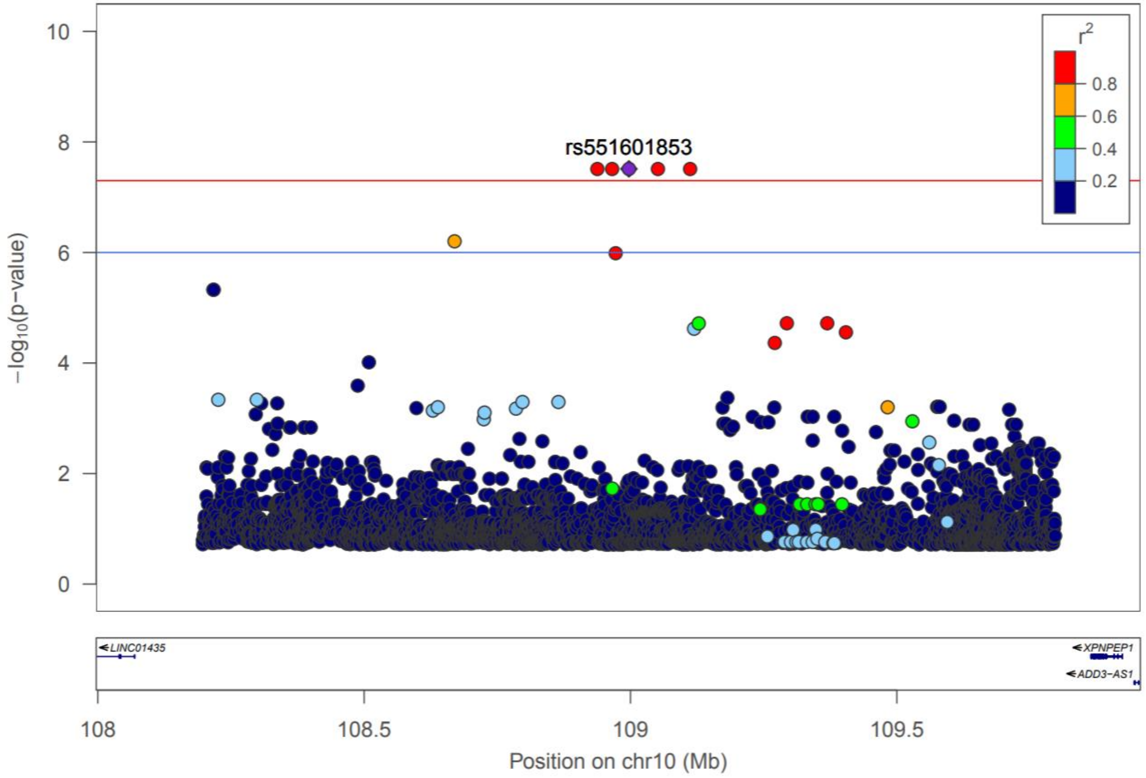


Single nucleotide variants are plotted with their P-values (-log10 values, left y-axis) as a function of build 38 genomic position on chromosome 10 (x-axis). Estimated recombination rates (right y-axis) are plotted to reflect the local linkage disequilibrium (LD) structure around the top associated single nucleotide variant rs551601853 (purple diamond) and correlated proxies (according to a blue to red scale from r2'=0 to 1). LD was calculated in non-diabetic African ancestry TOPMed cohorts.

**Supplementary Figure 5:** Regional HbA1c association plot in the *HK1* region (+/- 70kb around *HK1*).

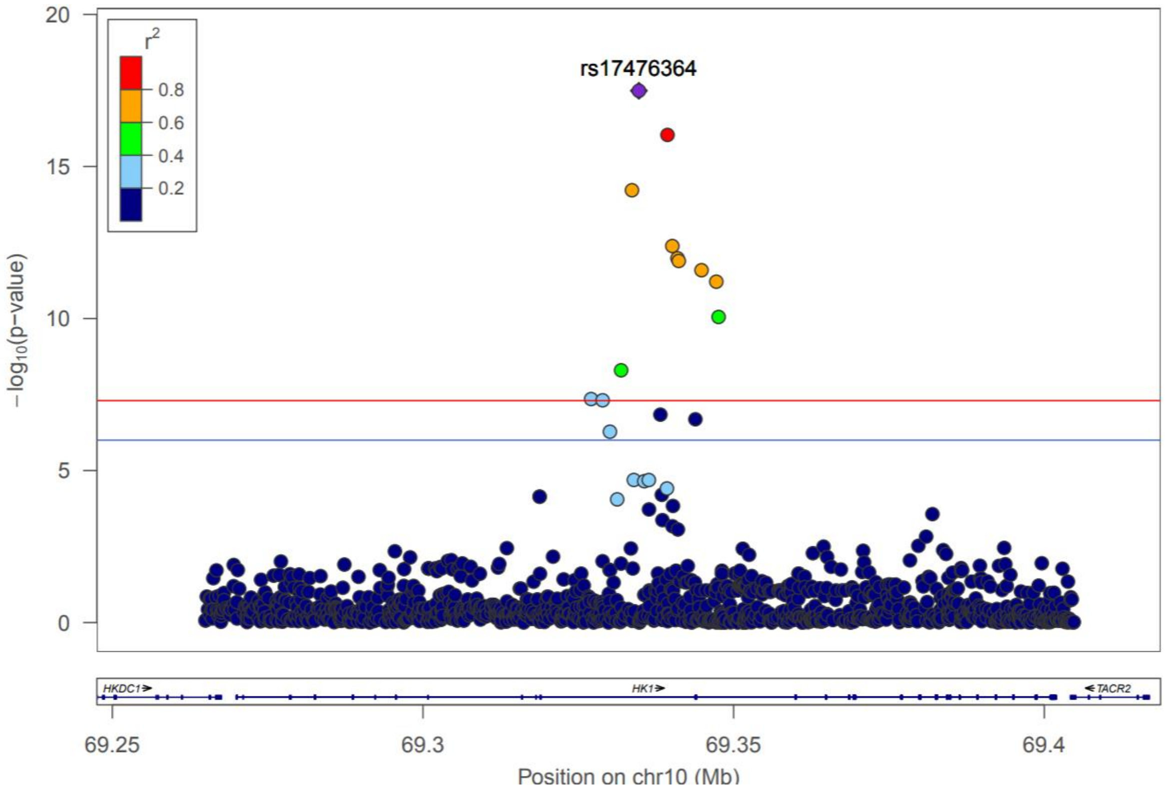


Single nucleotide variants are plotted with their P-values (-log10 values, left y-axis) as a function of build 38 genomic position on chromosome 10 (x-axis). Estimated recombination rates (right y-axis) are plotted to reflect the local linkage disequilibrium (LD) structure around the top associated single nucleotide variant rs17476364 (purple diamond) and correlated proxies (according to a blue to red scale from r2=0 to 1). LD was calculated in non-diabetic European ancestry TOPMed cohorts.

**Supplementary Figure 6:** Regional HbA1c association plot in the *G6PD* region (+/- 500kb around *G6PD*).

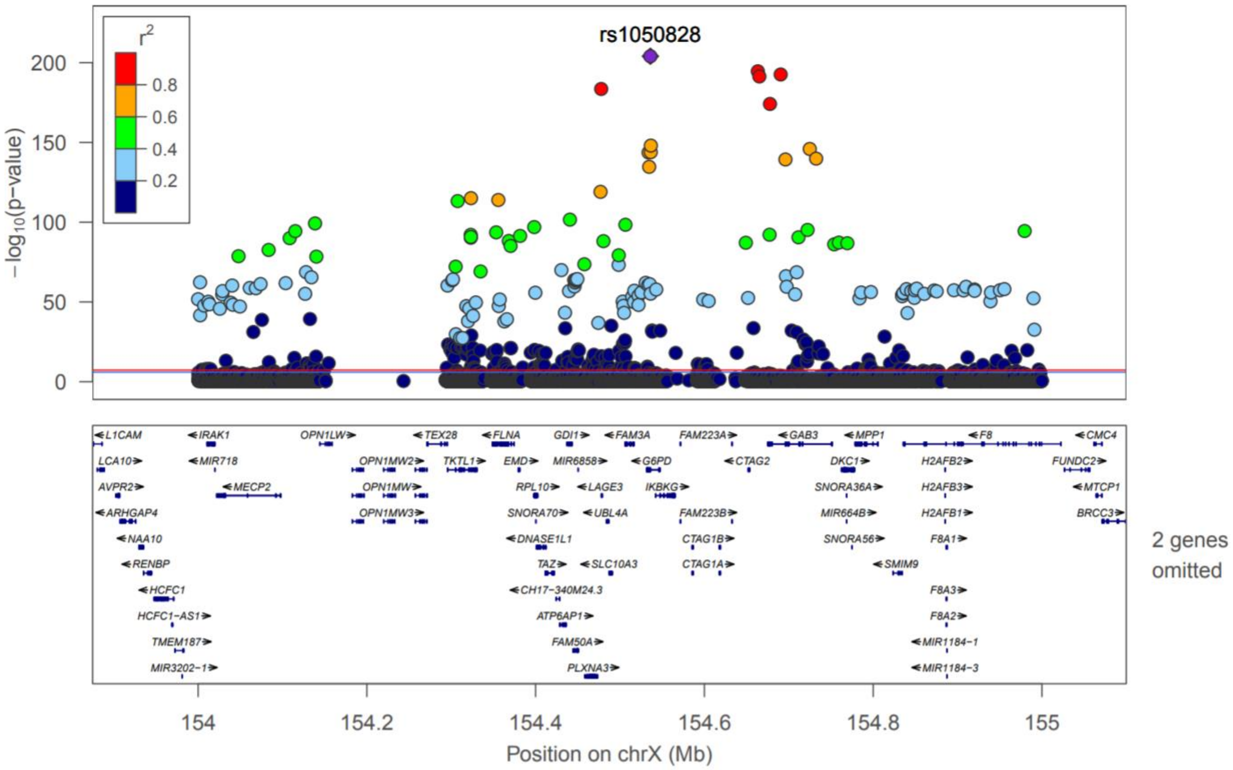


Single nucleotide variants are plotted with their P-values (-log10 values, left y-axis) as a function of build 38 genomic position on chromosome X (x-axis). Estimated recombination rates (right y-axis) are plotted to reflect the local linkage disequilibrium (LD) structure around the top associated single nucleotide variant rs1050828 (purple diamond) and correlated proxies (according to a blue to red scale from r2=0 to 1). LD was calculated in non-diabetic African ancestry TOPMed cohorts.
